# Transcriptomes of higher order thalamic nuclei in obsessive compulsive disorder

**DOI:** 10.1101/2025.04.22.650032

**Authors:** Shale A. Springer, Rishi Deshmukh, Lambertus Klei, Bernie Devlin, Jill R. Glausier, David A. Lewis, Susanne E. Ahmari

## Abstract

Obsessive-compulsive disorder (OCD) is a chronic psychiatric illness associated with altered function in cortico-striatal-thalamo-cortical (CSTC) circuits. In this pilot study, we examined differential RNA expression in the thalamus using postmortem human brain tissue samples from 11 subjects with OCD and 10 unaffected subjects. We individually dissected the mediodorsal magnocellular, mediodorsal parvocellular, and ventral anterior nuclei, which participate in orbitofrontal and anterior cingulate CSTC circuits most frequently associated with OCD, and the posterior ventrolateral nucleus, which participates in premotor and motor circuits that are increasingly implicated in OCD. Preselected GABAergic, glutamatergic and ion channel genes were analyzed via qPCR. Two genes required for GABA synthesis and release, *GAD1* and *SLC32A1*, were found to be downregulated in OCD subjects across all nuclei, and potassium channel *KCNN3* was upregulated. In parallel, we performed an exploratory total RNAseq differential expression analysis. We identified few (12 – 52) differentially expressed genes (DEGs) in each nucleus, and only one DEG in a pooled analysis of all nuclei. No DEGs were significant after correction for multiple comparisons. Investigation by model selection indicated that OCD diagnosis was not a useful factor in modelling gene expression in our dataset. OCD was also not associated with any modules of co-expressed genes identified using weighted gene correlation network analysis. Overall, we found minimal evidence of differential RNA expression in these thalamic nuclei in OCD. These findings contrast with our previous work including many of the same subjects where we found widespread differential mRNA expression in the orbitofrontal cortex and striatum in OCD.

## Introduction

Obsessive-compulsive disorder (OCD) is a chronic mental illness with a lifetime prevalence of 1-3% (Ruscio et al., 2010). People with OCD can experience significant impairment from the obsessions (recurring intrusive thoughts) and compulsions (repetitive behaviours, often in response to obsessions) characteristic of the disorder (Macy et al., 2013). Approximately 40-60% of OCD patients do not remit with first-line treatments, and 25% of those who do remit later relapse (Marcks et al., 2011; Sharma et al., 2014), highlighting the need for a better understanding of OCD pathophysiology to identify therapeutic targets.

Functional, structural, and molecular alterations in cortico-striatal-thalamo-cortical (CSTC) circuits have repeatedly been identified in subjects with OCD (Goodman et al., 2021; Shephard et al., 2021). In particular, the orbitofrontal cortex (OFC), anterior cingulate cortex (ACC), and striatum show hyperactivity and hyperconnectivity in patients with OCD at rest and during symptom provocation (Hazari et al., 2019; Norman et al., 2019; Perera et al., 2024). Although evidence for altered thalamic activity in OCD is inconsistent when analyzed as a single region (Hazari et al., 2019; Norman et al., 2019; Perera et al., 2024), medial and anterior thalamic regions do show altered functional connectivity with striatal, cortical and other thalamic regions in individuals with OCD (Gürsel et al., 2018; Jung et al., 2017; Kim et al., 2022). The anterior thalamus in particular serves as an functional connectivity “hub” central to OFC CSTC circuit activity in subjects with OCD, but not in unaffected subjects (Jung et al., 2017). In addition, OCD subjects show elevated choline levels indicative of inflammation or gliosis (Biria et al., 2021) in the medial thalamus.

In this study, we therefore selected individual nuclei within medial and anterior thalamic subregions that are connected to OFC- and ACC through CSTC circuits for post-mortem tissue analysis in subjects with and without OCD. The medial thalamus is primarily comprised of the mediodorsal nucleus (MD), a higher-order associative nucleus containing magnocellular (MDmc) and parvocellular (MDpc) subregions. MDmc primarily participates in limbic circuits involving the medial OFC and subgenual ACC, while MDpc is primarily engaged by associative circuits involving lateral OFC and dorsolateral PFC (Li et al., 2022; Mitchell and Chakraborty, 2013). In anterior thalamus, the ventral anterior nucleus (VA) is involved in limbic and supplemental motor circuits (McFarland and Haber, 2002, 2000; Rotge et al., 2008), and plays a role in compulsive and repetitive behaviors and Tourette’s Syndrome, which is highly comorbid with OCD (Huys et al., 2016; Rotge et al., 2012, 2008). We also identified a comparison nucleus outside of medial and anterior thalamus without strong OFC or ACC connection: the posterior ventrolateral nucleus (VLp). VLp participates in motor and premotor circuits, which have received increasing attention in OCD in recent years, and projects back to the striatum (de Wit et al., 2012; Khurshid, 2020; Lv et al., 2017; McFarland and Haber, 2002, 2000). Collectively, these four higher-order nuclei span limbic, associative, and motor CSTC circuits across the prefrontal cortex.

Analyzing individual thalamic nuclei requires a high level of anatomical resolution, such as that achievable through transcript expression analysis of postmortem human brain tissue. We have previously used qPCR and transcriptomic techniques to identify differential expression in the OFC, caudate, and nucleus accumbens, using tissue from a matched cohort of 8 OCD and 8 unaffected subjects (Piantadosi et al., 2021, 2019). In particular, we observed pronounced downregulation of glutamatergic signalling pathways and post-synaptic density genes, which have repeatedly been associated with OCD via familial analyses and near-significant genome-wide association studies (GWAS) hits (Bienvenu et al., 2009; Chakrabarty et al., 2005; Mahjani et al., 2021; Strom et al., 2024). Transcript expression profiles for striatal medium spiny neurons and cortical interneurons were also reduced in OCD subjects, while astrocyte and vascular cell expression profiles were elevated (Piantadosi et al., 2021). Differential transcript expression has also been identified in OFC, ACC, caudate, nucleus accumbens, and putamen tissue from a distinct OCD postmortem cohort of 6-8 subject pairs (de Oliveira et al., 2021; Lisboa et al., 2019). Epigenetic and transcriptomic data from this cohort were subsequently combined to generate networks of genes affected in OCD, which were enriched for synaptic, postsynaptic and immune system genes in all five regions (de Oliveira et al., 2021). In the present study, we extend these investigations to the thalamus using tissue from 11 subjects with OCD and 10 unaffected subjects. This expands on our previous cohort of 8 subject pairs, allowing us to assess differential expression of certain target genes across thalamic, cortical and striatal tissue samples from the same subjects. We here assess differential transcript expression via 1) targeted qPCR analysis of genes associated with OCD, 2) exploratory total RNAseq differential expression analysis, and 3) weighted gene correlation network analysis (WGCNA) of transcriptomic data.

Our targeted qPCR analysis focused on glutamatergic and GABAergic synaptic genes and ion channels involved in thalamocortical cell burst firing, a phenomenon closely linked to the excitatory/inhibitory balance in the thalamus. We selected nine glutamatergic synapse genes that are essential to glutamatergic function and/or have been associated with OCD and compulsive behavior, including five genes downregulated in OFC and striatal tissue of our overlapping postmortem cohort (Bienvenu et al., 2009; Chakrabarty et al., 2005; de Oliveira et al., 2021; Grünblatt, 2021; Lisboa et al., 2019; Piantadosi et al., 2021, 2019). GABAergic target genes identified were the vesicular inhibitory amino acid transporter, GABA synthesis enzymes glutamate decarboxylase 1 and 2, receptor scaffolding gene *GABARAP*, and two extrasynaptic ionotropic GABA receptor subunits. These extrasynaptic GABA receptors mediate tonic inhibition levels in the thalamus and thereby regulate thalamocortical cell firing mode (Cope et al., 2005; Paydar et al., 2014), which has been proposed to underlie altered thalamic oscillations in Tourette’s syndrome (Bour et al., 2015; Priori et al., 2013; Shao et al., 2021). Similar alterations have also been identified in subjects with OCD (Li et al., 2019; Perera et al., 2023). We anticipated that cortical and striatal alterations previously documented in OCD were likely to affect E/I balance and therefore burst firing regulation in these thalamic nuclei; we therefore selected five cation channels involved in execution or regulation of thalamocortical cell burst firing for targeted analysis (Jin et al., 2000; Kasten et al., 2019; Kim et al., 2001; Shao et al., 2021; Zobeiri et al., 2019).

In parallel with our targeted analyses, we performed total RNA sequencing followed by transcriptomic differential expression and co-expression network analyses. These untargeted exploratory analyses have the potential to identify novel pathways of interest. In cortical and striatal tissue, similar methods did identify differential expression of vascular and angiogenic pathway genes not previously associated with OCD (de Oliveira et al., 2021; Lisboa et al., 2019; Piantadosi et al., 2021). Finally, we used WGCNA to identify modules of co-expressed genes in our transcriptomic dataset, and compared module expression values between OCD and unaffected subjects. Such modules typically share pathway membership and functional relevance and can identify more coordinated, subtle shifts in transcript expression (Langfelder and Horvath, 2008; Zhang and Horvath, 2005). Using these complementary techniques we sought to investigate the effect of OCD on transcript expression in individual thalamic nuclei.

## Materials and Methods

### Postmortem Human Subjects

Brain tissue from 22 postmortem human subjects was obtained through the University of Pittsburgh Brain Tissue Donation Program. Tissue was acquired during autopsies conducted by the Allegheny County Medical Examiner’s Office (Pittsburgh, PA) with the consent of next-of-kin. DSM-IV diagnoses were made by the consensus of an independent committee of experienced clinicians, who conducted structured interviews with family members and/or reviewed the subjects’ clinical records. The same process was used to determine unaffected comparison subjects to be free of any lifetime psychiatric illnesses. Each subject with OCD was matched to an unaffected comparison subject for sex and as closely as possible for age, postmortem interval (PMI), brain pH, and tissue storage time (Supplemental Table 1) to reduce biological variance between groups. One unaffected subject (pair 8) was excluded from analysis due to poor-quality expression data (see RNAseq methods) and has been excluded from cohort summary characteristics. Unaffected subjects had significantly longer tissue storage time at −80°C (Table 1). Groups did not differ in other tissue or subject characteristics. All procedures were approved by the University of Pittsburgh’s Committee for the Oversight of Research and Clinical Training Involving Decedents and Institutional Review Board for Biomedical Research.

**Table 1:** Summary subject characteristics.

|  | Unaffected Subjects<br>(n = 10) | OCD Subjects<br>(n = 11) | F | p |
| --- | --- | --- | --- | --- |
| Sex (n) | 4M/6F | 5M/6F |  |  |
| Age (years) | 44.8 (±11.8) | 45.2 (±12.6) | 0.005 | 0.944 |
| Body Mass Index (BMI) | 31.0 (±7.0) | 28.2 (±6.7) | 0.832 | 0.373 |
| Postmortem interval (hours) | 16.1 (±5.0) | 17.5 (±7.5) | 0.254 | 0.620 |
| Brain pH | 6.6 (±0.2) | 6.6 (±0.2) | 0.455 | 0.509 |
| Subject RIN | 7.8 (±0.6) | 7.9 (±0.6) | 0.159 | 0.695 |
| Tissue storage time (months) | 157.9 (±48.9) | 102.7 (±43.0) | 7.580 | **0.013\*** |
| Suicide (n) | 0 | 3 |  |  |
| Antidepressant use ATOD (n) | 0 | 8 |  |  |
| Number of subjects (n) or group mean (+/- SD) values are reported. Abbreviations: ATOD = At time of death; RIN = RNA Integrity Number |  |  |  |  |

### Tissue Collection

MDmc, MDpc, VA, and VLp nuclei were collected via cryostat from coronal blocks of the right hemisphere. Collection regions were identified by anatomical landmarks with reference to Atlas of the Human Brain (4^th^ Edition) (Mai et al., 2015). Target collection areas are shown in Supplemental Figure 1.

### Quantitative PCR

Quantitative PCR data is reported from 10 OCD-unaffected subject pairs for MDpc and VA and from 8 pairs for MDmc and VLp (Supplemental Table 3), excluding any samples that did not yield adequate RNA or failed to amplify housekeeping genes. We also previously collected expression data from OFC and striatal regions for pairs 1-8, allowing us to compare differential expression across multiple CSTC regions within the same subjects (Piantadosi et al., 2019). Cross-region data was available for 7 pairs after the exclusion of pair 8 from thalamic analyses. Using this data, we performed parallel differential expression analyses of thalamic and OFC/striatal data for eight target genes.

RNA was extracted using a RNeasy Plus Micro kit (QIAGEN, Valencia, CA). RNA integrity number (RIN) and concentration were assessed using the Agilent 2100 Bioanalyzer (Agilent Technologies, Santa Clara, CA). Complementary DNA (cDNA) was generated from RNA using the qScript cDNA synthesis kit (QuantaBio, Beverly, MA). All primer sequences are listed in Supplemental Table 2. Paired samples were run on the same qPCR plates to account for inter-run variability. Samples were run in technical triplicate. Transcript data was excluded for individual samples if the standard error of the cycle threshold (CT) values for its technical replicates was greater than 0.3 cycles. mRNA expression levels were normalized by subtracting the mean CT of the target gene from the mean CT of two housekeeping genes [Actin (*ACTB*) and β-2 microglobulin (*B2M*)] and calculating the geomean of these delta CT (dCT) values. Because CT values are points on a logarithmic curve, expression levels are reported as 2^-dCT^.

Individual general linear models were created to model expression for each gene using the *lme4* package in R. Diagnosis, brain region, and diagnosis by region interaction were included as fixed effects. Subject ID was included as a random effect. A stepwise selection procedure was then run to select additional covariates that improved fit for individual models, similar to previously described procedures (Fromer et al., 2016; Piantadosi et al., 2021). Additional covariates were selected for *GAD1* (suicide) and *GABRD* (pH). For the cross-region analysis of the first 7 subject pairs, suicide was a covariate for *GAD1* and *GAD2* in thalamic models, and no covariates were selected for cortical and striatal models. Significance tests were generated using the R package *car*. Adjustment for multiple comparisons across 20 genes was performed using the Benjamini-Hochberg (BH) method and FDR = 0.05. Uncorrected p values are reported in the text; corrected values are reported in Supplemental Table 4. Following significant diagnosis by region interactions, post-hoc paired t-tests were performed using the R package *stats* and corrected for multiple comparisons across four regions.

### RNA Sequencing

cDNA libraries were generated using Illumina TruSeq Stranded Total RNA kit for 86 samples (RNA was not available for two VA samples). Mean sample RIN was 7.7 (range 6.0 to 9.4) (Supplemental Table 5). Paired-end (50bp x2) total RNA sequencing was performed using S2 flow cells on the Illumina NovaSeq 6000 platform at a target depth of 40 million reads per sample. Reads were aligned to human genome reference sequence GRCh38.p13 and annotated with Ensemble Archive Release v106 using CLC Genomics Workbench 22 (QIAGEN). Over 90% of reads were mapped for each sample, with 83-95% mapped in pairs (Supplemental Figure 2). One sample was excluded due to low total read count, and three outliers were identified and removed during data analysis (Supplemental Figure 2). The sole remaining sample from the unaffected subject in pair 8 was also removed, leaving samples from 11 OCD subjects and 8-10 unaffected comparison subjects per region. Of 37,704 protein coding or long noncoding RNA (lncRNA) genes identified, 14,517 met our expression threshold and passed QC, defined as at least 1 count per million in all regions and with coefficient of variance < 3 across all samples. Differential expression was assessed using generalized linear regression. Covariates were selected following a stepwise forward procedure as previously described (Fromer et al., 2016; Piantadosi et al., 2021) beginning with a model including diagnosis as the only predictor. Each covariate was assessed for improved model fit as assessed by Bayesian information criteria (BIC) compared to the null model, and the covariate that improved fit for the largest number of genes was added to the model. This was repeated until no covariate improved fit for ≥5% of genes. Following this step, surrogate variable analysis (SVA) was performed using the R package *sva* (Leek et al., 2012). For global thalamic expression, the selected model included region, pH, and 3 surrogate variables (SV) in addition to diagnosis. For regional models the following additional covariates were selected: MDmc: age and three SVs; MDpc: pH and one SV; VA: two SVs; VLp: pH, RIN and two SVs. Estimated voom precision weights were calculated for each gene and applied to weighted linear regression models fit using limma (Abdul et al., 2017; Law et al., 2014; Ritchie et al., 2015). For each gene, significance from the case-control contrast is evaluated as a limma-moderated t-test of the effect of diagnosis in the context of the linear model. Adjustment for multiple testing was performed using the BH procedure.

### WGCNA

Weighted gene correlation network analysis (WGCNA) was conducted to identify modules of co-expressed genes in RNAseq expression data (Langfelder and Horvath, 2008; Zhang and Horvath, 2005). 5000 genes with highest variance across all RNAseq samples were selected for WGCNA, including 3,966 protein-coding genes and 1,034 lncRNA genes. Signed-hybrid weighted co-expression networks were constructed for each region using data from both OCD and unaffected subjects (19-21 samples). Since this is the recommended minimum of 20 samples, biweight midcorrelation was used to build more robust networks (Ballouz et al., 2015). Soft thresholding power was selected for each region at a scale-free topology fit index threshold of 0.8. Selected powers were 5, 4, 5, and 4 for MDmc, MDpc, VA, and VLp respectively. Non-overlapping modules of co-expressed genes were identified by dynamic branch cutting of hierarchically clustered dendrograms (Langfelder et al., 2008), with a minimum of 60 genes per module. We used hierarchical clustering of module eigengenes, defined as the first principal component for each module (Langfelder and Horvath, 2007), to merge modules with dissimilarity less than 0.25. Module color names are randomly assigned within each region. Over-representation analysis was performed using WebGestalt in R (Liao et al., 2019), and genes that did not map to any GO term (n = 1146, 88% lncRNA) were automatically excluded.

Module-trait relationships were assessed by correlating module eigengene values with sample/subject traits that were factors of interest (OCD or depression diagnosis); were included in one or more RNAseq expression model(s); or differed between OCD and unaffected groups. Correlation significance was assessed using a Student’s asymptotic t-test followed by BH correction for multiple comparisons across the number of traits tested.

## Results

### Target gene expression

Twenty target genes were included in qPCR analyses. A mixed linear model was generated to evaluate expression of each gene, including OCD diagnosis and thalamic region as fixed effects and pair as a random effect. Additional tissue or subject covariates were included if they met criteria via a stepwise forward selection process. Significance is reported at unadjusted p < 0.05 based on *a priori* gene selection.

A main effect of OCD diagnosis was observed for *GAD1* (F_1,62_ = 8.16, p = 0.006) and VIAAT gene *SLC32A1* (F_1,59_ = 4.63, p = 0.036). Both genes showed lower mRNA expression in OCD subjects (Figure 1a,b). Death by suicide improved fit for our model of *GAD1* expression and showed a significant main effect (p = 0.0001); within-group comparison revealed higher *GAD1* expression in OCD subjects who died by suicide (n = 3) than those who did not (n = 8) (t_10_ = 2.71, p = 0.02). *GAD2* showed no main or interaction effect of OCD diagnosis (Supplemental Figure 3) and no additional covariates. OCD had no significant effect on post-synaptic GABAergic genes *GABARAP*, *GABRD* or *GABRA4* (Supplemental Table 4a; Supplemental Figure 3). All GABAergic genes showed a main effect of region (no diagnosis by region interaction), with presynaptic GABAergic genes most highly expressed in MDpc.

**Figure 1.**
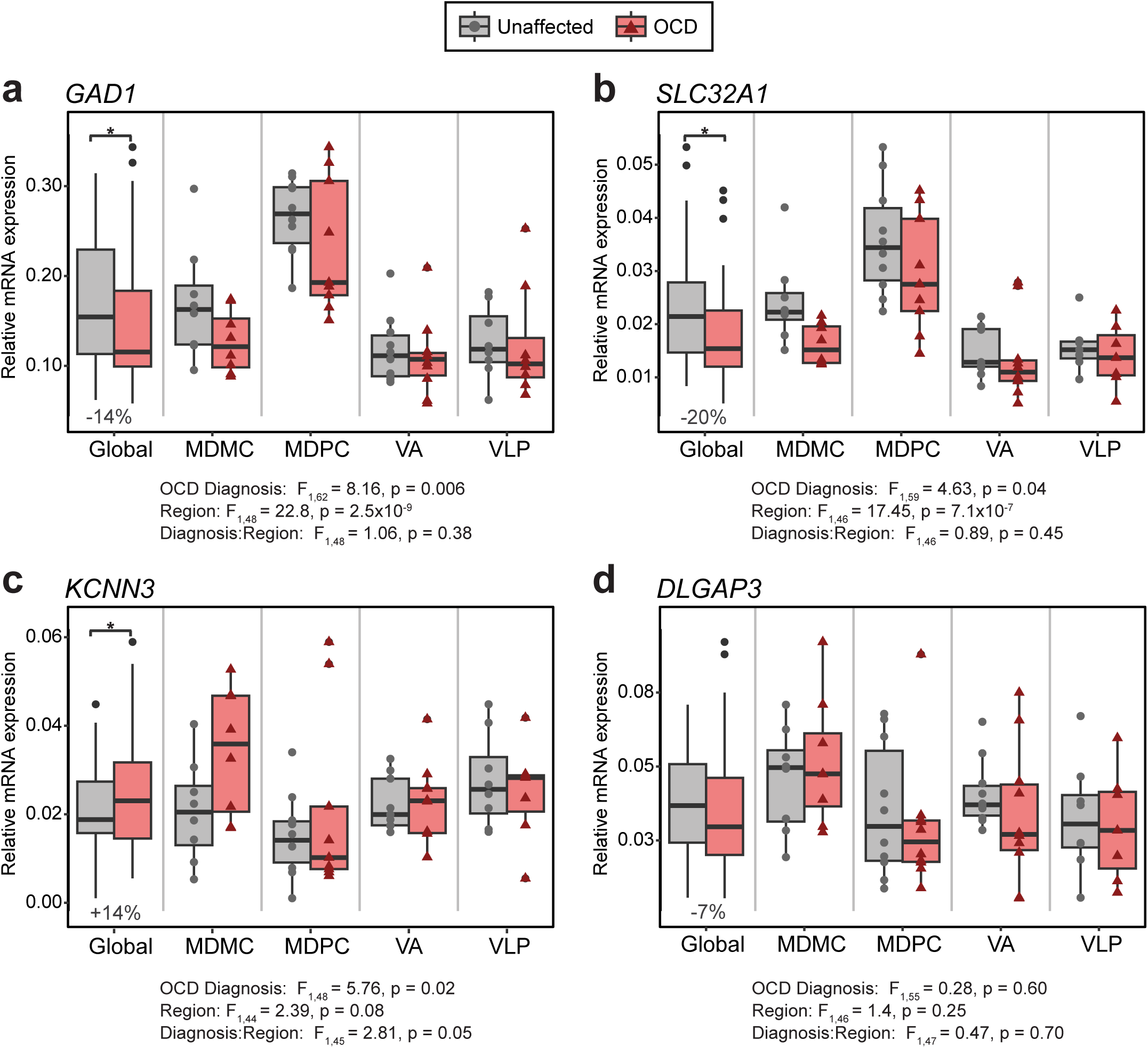
Differential expression of selected OCD target genes between OCD and unaffected subjects evaluated using qPCR. Boxplots depict relative expression levels of (a) GAD1, (b) SLC32A1, (c) KCNN3, and (d) DLGAP3 in each thalamic nucleus and across all thalamic samples (Global). *p < 0.05 for main effect of diagnosis

Expression of SK-type potassium channel *KCNN3* was elevated in OCD subjects (F_1,48_ = 5.76, p = 0.02) with a trend towards a diagnosis by region interaction (F_3,45_ = 2.81, p = 0.05). No effect of OCD diagnosis was observed for potassium channels *KCNA2* or *KCNMA1*, cation channel *HCN4*, or low voltage gated calcium channel *CACNA1G* (Supplemental Figure 3, Supplemental Table 4a). We also found no effect of OCD diagnosis on genes related to glutamatergic signalling and synaptic structure, including post-synaptic scaffolding genes *DLGAP3* (F_1,55_ = 0.28, p = 0.60) (Figure 1d) and *DLGAP4* (F_1,50_ = 0.02, p = 0.89), NMDA-type ionotropic glutamate receptor subunit *GRIN2B* (F_1,54_ = 0.07, p = 0.79), and presynaptic glutamate transporter *SLC1A1* (F_1,59_ = 0.03, p = 0.87) (Supplemental Figure 3). Ionotropic glutamate receptor subunits *GRIA4* and *GRIN1*, metabotropic glutamate receptors *GRM3* and *GRM5*, and vesicular glutamate transporter *SLC17A6* were also unaffected by OCD diagnosis (Supplemental Figure 3, Supplemental Table 4a).

We previously identified differential mRNA expression in two medial OFC regions (BA11 and BA47m), caudate, and nucleus accumbens tissue from an overlapping postmortem OCD cohort (Piantadosi et al., 2019). Four GABAergic (*GAD1, GAD2, SLC32A1*, and *GABARAP*) and four glutamatergic (*DLGAP3*, *DLGAP4, GRIN2B*, and *SLC1A1*) genes were also assessed in these regions. In order to compare results across regions, we analyzed data from seven subject pairs included in both studies (Supplemental Table 4b). *GAD1* and *SLC32A1* were significantly downregulated in the thalamus and showed a significant diagnosis by region interaction in mOFC and striatum, but post-hoc testing in cortex and striatum found no significant group differences (Table 2; Supplemental Table 4c). Consistent with previous results, we found a strong downward effect of OCD on *GRIN2B, SLC1A1*, and *DLGAP3* expression in mOFC and striatum, but not in thalamus (Table 2). *GAD2*, *GABARAP* and *DLGAP4* were not differentially expressed in either region, though *DLGAP4* showed a trend towards downregulation in OCD in cortex and striatum (F_1,30_ = 3.76, p = 0.06) (Supplemental Table 4b).

**Table 2:** Differential expression of select genes in thalamus, mOFC and striatum.

| Gene | Regions | Factor | F | df | Residual df | p |
| --- | --- | --- | --- | --- | --- | --- |
| *GAD1* | Thalamus | OCD | 11.13 | 1 | 39 | **0.002\*** |
|  |  | OCD:Region | 0.85 | 3 | 31 | 0.48 |
|  | mOFC, Caudate, NAc | OCD | 0.13 | 1 | 37 | 0.72 |
|  |  | OCD:Region | 4.82 | 3 | 36 | **0.006\*** |
| *SLC32A1* | Thalamus | OCD | 5.70 | 1 | 34 | **0.023\*** |
|  |  | OCD:Region | 1.18 | 3 | 30 | 0.33 |
|  | mOFC, Caudate, NAc | OCD | 0.005 | 1 | 40 | 0.95 |
|  |  | OCD:Region | 4.86 | 3 | 36 | **0.006\*** |
| *GRIN2B* | Thalamus | OCD | 0.08 | 1 | 31 | 0.78 |
|  |  | OCD:Region | 0.63 | 3 | 31 | 0.60 |
|  | mOFC, Caudate, NAc | OCD | 6.86 | 1 | 34 | **0.013\*** |
|  |  | OCD:Region | 0.89 | 3 | 36 | 0.46 |
| *DLGAP3* | Thalamus | OCD | 0.0002 | 1 | 33 | 0.99 |
|  |  | OCD:Region | 0.80 | 3 | 31 | 0.50 |
|  | mOFC, Caudate, NAc | OCD | 11.37 | 1 | 48 | **0.002\*** |
|  |  | OCD:Region | 0.69 | 3 | 36 | 0.57 |
| *SLC1A1* | Thalamus | OCD | 0.002 | 1 | 32 | 0.96 |
|  |  | OCD:Region | 2.12 | 3 | 30 | 0.12 |
|  | mOFC, Caudate, NAc | OCD | 15.77 | 1 | 48 | **0.0002\*** |
|  |  | OCD:Region | 2.67 | 3 | 36 | 0.06 |

### Transcriptome differential expression

In transcriptomic analyses, gene expression was modelled by OCD diagnosis with additional covariates chosen via stepwise forward selection, as previously described (Piantadosi et al., 2021) (Supplemental Table 6a). Subject pair was included as a random effect, and supplemental variable analysis (SVA) was performed for each contrast to improve model fit. Regions were analyzed individually and pooled for a “global thalamic” analysis. Hierarchical sample clustering indicated moderate clustering by region but not by diagnosis (Supplemental Figure 4).

No DEGs were identified in the global thalamic analysis at FDR-adjusted p < 0.05 (Figure 2a; Supplemental Table 7a). The distribution of unadjusted p values was skewed left, indicating a preferential failure to reject the null hypothesis (Figure 2b). Only one DEG (antisense RNA *L3MBTL2-AS1*) was identified at a threshold of unadjusted p < 0.01 and absolute fold change >20%. No qPCR target genes were differentially expressed, including *GAD1* (log_2_FC = −0.03, p = 0.71), *SLC32A1* (log_2_FC = −0.13, p = 0.30) and *KCNN3* (log_2_FC = −0.03, p = 0.74). Within-subject RNAseq and qPCR expression values were significantly correlated for 10 target genes, including *GAD1* and *SLC32A1* but not *KCNN3*, and 15 of the 20 target genes showed a consistent direction of change across methods (Supplemental Table 4d).

**Figure 2.**
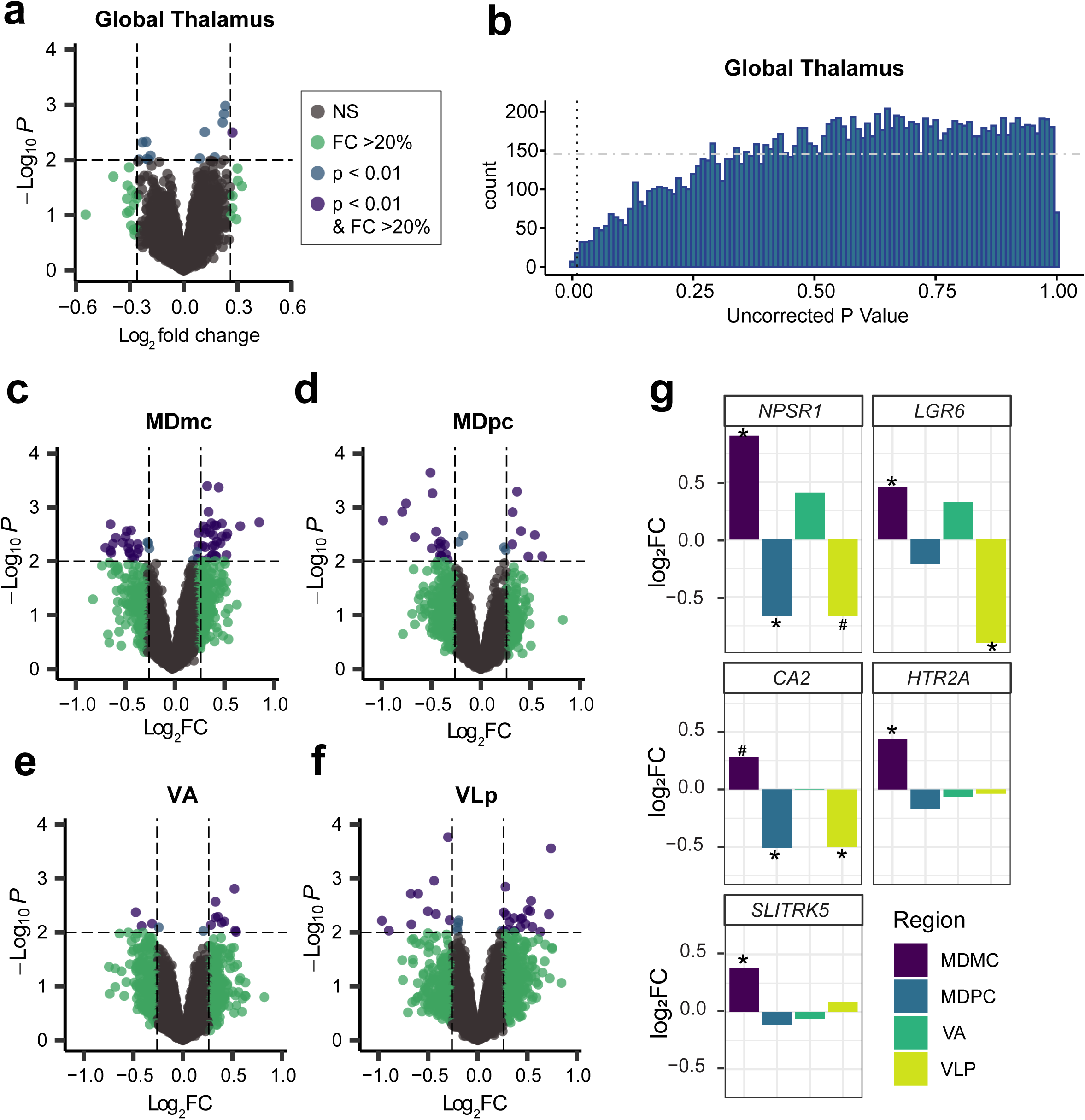
Differential transcript expression between subjects with and without OCD evaluated using total RNA sequencing. (a) Volcano plot depicts differential expression results for all transcripts (n = 14,517) across all thalamic nuclei (global thalamic analysis). (b) Distribution of uncorrected p-values for OCD differential expression contrasts in the global thalamic analysis. A significance cutoff of p = 0.01 is indicated by the vertical dotted line. The expected distribution of p-values distributed at random (i.e. expected distribution under the null hypothesis) is indicated by the horizontal dash-dot line. Observed leftward skew indicates fewer significant contrasts than expected by chance. (c-f) Volcano plots of differential RNA expression results from (c) mediodorsal magnocellular (MDmc), (d) mediodorsal parvocellular (MDpc), (e) ventral anterior (VA) and (f) ventrolateral posterior (VLp) nuclei. (g) Bar plot indicates the log_2_FC between OCD and unaffected subjects for select differentially expressed genes (DEGs) in each thalamic nucleus. *p < 0.01 and FC >20%. #p < 0.05 and FC > 20%.

Regional RNAseq analyses of MDmc, MDpc, VA and VLp yielded 52, 24, 12 and 30 DEGs respectively at a threshold of unadjusted p < 0.01 and absolute fold change > 20% (Figure 2c-f, Supplemental Table 7b-e). Two OCD-associated synaptic genes were DE only in MDmc, namely *HT2RA* (log_2_FC = 0.44, p = 0.005) and *SLITRK5* (log_2_FC = 0.38, p = 0.001); (Figure 2g). Three genes-*NPSR1*, *LGR6* and *CA2–* showed substantial DE in multiple regions, consistent with regional differences in OCD-related differential expression. Each of these three genes were upregulated in MDmc and downregulated in VLp and MDpc (Figure 2g). All three showed a positive fold-change in VA. However, as in the global analysis, no DEGs were significant after correcting for multiple comparisons (Supplemental Figure 5).

As our primary factor of interest, OCD diagnosis had been included in all RNAseq linear regression models prior to covariate selection. However, the lack of differential expression and evidence of model overfit led us to repeat stepwise covariate selection from a null model including only the intercept, with no factors of interest. We found that OCD diagnosis improved model fit vs. the intercept model for only two genes (0.01%) in the global thalamic analysis and 24 – 55 genes (0.2% – 0.4%) in each regional analysis (Supplemental Table 6b). OCD diagnosis did not meet our threshold for covariate inclusion, i.e. improved model fit for ≥5% of genes in the analysis. For comparison, we performed an exploratory analysis of differential expression between thalamic regions, since this variable did improve model fit for >5% of genes. We were able to identify differential expression at FDR-corrected p < 0.05, ranging from 2 DEGs for MDmc vs. VA to 4,315 DEGs for MDpc vs. VLp. All 6 between-region contrasts showed typical right-skewed p-value distributions (Supplemental Figure 6). The bias towards the null observed in OCD differential expression analysis likely originates from model overfitting, as OCD diagnosis was not a useful factor for modelling gene expression in our dataset.

### Weighted gene co-expression network analysis

We generated a weighted gene correlation network for each region using the 5,000 genes with highest variance across the entire dataset. Networks were based on data from both OCD and unaffected subjects. We identified 11 or 12 modules (depending on the region being examined) of co-expressed genes within each region, which are identified by arbitrary color names (Figure 3, Supplemental Table 8). Two modules in each network showed significant overrepresentation of synapse-related genes, with one primarily enriched for post-synaptic and the other for pre- or trans-synaptic GO terms (Supplemental Table 9). These modules are annotated “P” or “S” respectively in Figure 3. We then performed module-trait correlation analyses for OCD diagnosis, depressive disorder diagnosis (n = 7 subjects), and covariates that influenced our RNA expression models (age, pH, and RIN) or differed between subject groups (tissue storage time). OCD diagnosis was not correlated with module eigengene values (MEs) for any module (Figure 3; Supplemental Table 10). Depressive disorder diagnosis showed a nominally significant negative correlation with Mes for *MDpc-yellow* (R^2^ = −0.50, p = 0.03) and *VLp-yellow* (R^2^ = −0.46, p = 0.04), which were both enriched for immune response-related GO terms (Supplemental Table 9). However, these correlations did not survive correction for multiple comparisons across the number of modules in each region. Significant module-trait correlations were also identified for age, pH, and subject RIN, but not tissue storage time (Figure 3, Supplemental Table 10).

**Figure 3.**
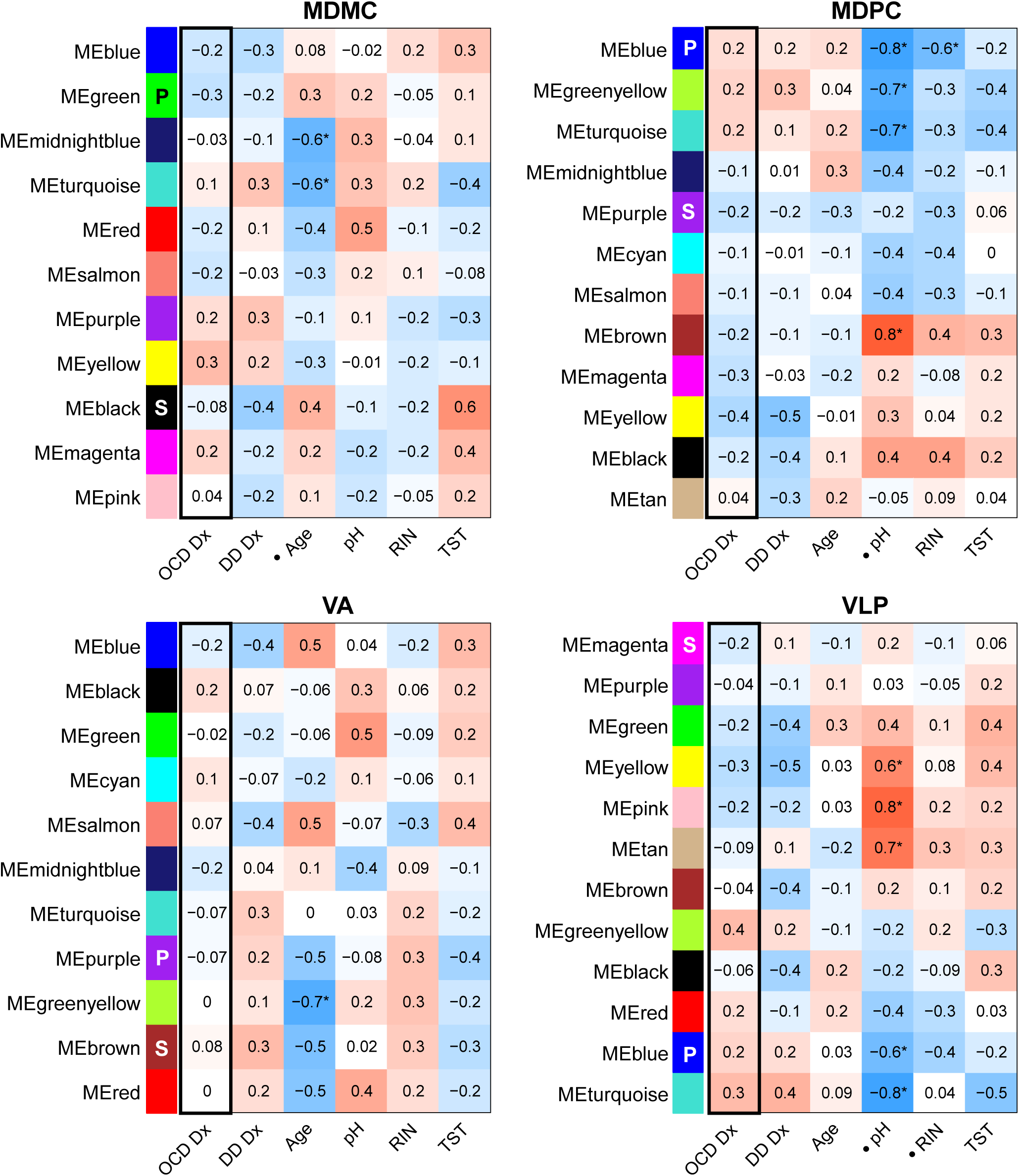
WGCNA module-trait correlation. Correlation (R2 value) between WGCNA module eigengene values for each region and sample/subject traits. Modules are identified by arbitrary color names within each region. Modules enriched for GO:0045202 (“synapse”) or GO:0014069 (“postsynaptic density”) are marked with S or P respectively. 1 indicates a trait that met criteria for inclusion in the region’s RNAseq expression model. For OCD and depressive disorder diagnoses, 0 = no diagnosis and 1 = diagnosis. DD Dx = Depressive disorder diagnosis; RIN = RNA integrity number; TST = tissue storage time. *FDR-corrected p < 0.05.

## Discussion

In this study, we evaluated the relationship between OCD diagnosis and RNA expression in the thalamus using targeted qPCR analysis and total RNA sequencing in a small pilot cohort. We selected four thalamic nuclei based on their participation in OCD-related CSTC circuits: MDmc, MDpc, VA, and VLp. Using a targeted qPCR approach, we observed downregulation of *GAD1* and *SLC32A1* and upregulation of potassium channel *KCNN3* across all four nuclei. Glutamatergic synapse genes were not differentially expressed. In our wider transcriptomic analysis, no DEGs were significant after FDR correction and a bias towards the null was observed among uncorrected p-values. Furthermore, OCD diagnosis did not improve gene expression model fit for the majority of genes analysed. We also found no relationship between OCD diagnosis and any WGCNA modules.

Two DEGs identified via qPCR, *GAD1* and *SLC32A1*, are involved in GABA synthesis and release. The GABAergic thalamic interneurons (TINs) that express these genes enable communication between thalamocortical cells, which do not form direct synapses with one another (Sherman, 2004). In either case, local circuit alterations are likely to alter integration, processing, and coordination between thalamocortical cells. Using data from the first seven subject pairs, we also found that *GAD1* and *SLC32A1* were downregulated in the thalamus but not the OFC or striatum. Reduced expression of GABA release machinery in TINs could indicate a compensatory response to altered circuit activity, such as increased inhibitory input from the basal ganglia, or could itself contribute to hyperactivity in this circuit. However, differential expression of target GABAergic genes in OCD has not been consistent between qPCR and RNAseq methods. *SLC32A1* and *GAD1* downregulation in the thalamus were not confirmed by transcriptomic analysis. In the OFC and striatum, *SLC32A1* was found to be downregulated in the transcriptomic analysis but not targeted qPCR (Piantadosi et al., 2021, 2019). These preliminary findings should therefore be interpreted with caution, though they encourage future investigation of GABAergic signaling alterations in OCD.

Our interpretation of *KCNN3* upregulation in OCD subjects faces similar limitations. *KCNN3* encodes the calcium-activated potassium channel SK3, which contributes to afterhyperpolarization and decreases tonic firing rate and gain in thalamocortical neurons (Kasten et al., 2007). Its upregulation could indicate conditions conducive to burst firing (Shao et al., 2021). Alternatively, *KCNN3* upregulation could indicate response to inflammation or gliosis, which are elevated in the medial thalamus in OCD (Biria et al., 2021; Dolga et al., 2012; Gerentes et al., 2019; Siddiqui et al., 2014). However, other genes involved in these coordinated processes were not differentially expressed, and *KCNN3* elevation was not confirmed in transcriptomic analyses.

Differential expression of glutamatergic and PSD genes was absent in both qPCR and RNAseq analyses. In fact, no genes were differentially expressed in thalamic transcriptomic analyses after FDR correction, and relatively few nominally significant DEGs were identified at a more lenient threshold. We failed to reject the null hypothesis more often than would be expected by chance. Surrogate variable analysis to account for unknown drivers of variance did not eliminate this bias. Given the lack of support for our initial hypothesis, we reassessed the role of OCD in our models of gene expression. We found that OCD diagnosis improved model fit for less than 0.4% of genes, indicating that it was not a useful factor for modelling gene expression in our dataset.

Although the relationship between OCD and gene expression in the thalamus appeared minimal overall, the lack of consensus between individual thalamic nuclei could inform future analyses. We identified 12 – 52 DEGs in each nucleus and only one DEG in a pooled analysis of all samples, despite the relatively low power of individual regional analyses. Three DEGs were identified in multiple nuclei, and all three were upregulated in MDmc and downregulated in MDpc and/or VLp. This does not reflect underlying regional expression patterns in which MDmc and MDpc samples were most similar (Supplemental Figure 2). We previously observed that pooling OFC and striatal samples increased power to identify OCD-related DEGs overall, despite dissimilarity between the regions at baseline, due to the presence of concordant differential expression (Piantadosi et al., 2021). Future studies should consider the possibility that OCD affects individual thalamic nuclei in distinct ways despite the lack of a clear effect of OCD on the region as a whole.

Our findings are limited by the small and heterogenous nature of our subject cohort, and the lack of an effect of OCD diagnosis in our dataset does not rule out OCD-related differential expression in the thalamus altogether. We were not powered to assess sex differences in this cohort, and were unable to test the effect of comorbid disorders or medication use beyond testing whether MDD diagnosis, antidepressant use at time of death, or suicide were useful covariates. Tissue storage time also significantly differed between diagnostic groups, but it was not included in our models because it was confounded with diagnosis. Given the limited diagnostic differences we observed, our results are consistent with the other findings that flash-frozen brain tissue can be stored at −80°C for decades with only subtle effects on RNA integrity (White et al., 2018). Furthermore, we have previously been able to detect substantial differential expression in OFC and striatal tissue from a cohort of 8 matched pairs, including the majority of subjects in the present study, with many of the same limitations (Piantadosi et al., 2021, 2019).

The relative absence of OCD-related differential expression in the thalamus supports a preferential vulnerability of corticostriatal synapses in OCD relative to thalamocortical, thalamostriatal, or corticothalamic synapses. Differential expression in cortical and striatal regions has been identified in two distinct postmortem cohorts, each including only 6-8 OCD and unaffected subject pairs and one largely overlapping with the present study (de Oliveira et al., 2021; Lisboa et al., 2019; Piantadosi et al., 2021). Our parallel analyses of qPCR data from 7 subject pairs allowed us to compare these regions more directly, showing downregulation of glutamatergic and PSD scaffolding genes only in the OFC and striatum. This is consistent with the current focus of the field, which has been motivated by substantial evidence of cortical and striatal dysfunction in OCD and preclinical models of compulsive behavior (Shephard et al., 2021; Shmelkov et al., 2010; Welch et al., 2007). Our findings support the interpretation that thalamic synaptic gene expression is relatively preserved in OCD within the same CSTC circuits where corticostriatal synaptic deficits have been reported, including in these same individuals. In addition to continued focus on corticostriatal synapses, future research should consider other elements of these CSTC circuits, such as the output nuclei of the basal ganglia, the subthalamic nucleus (STN), and the thalamic reticular nucleus, which could mitigate the impact of cortical and striatal dysfunction on the thalamus in OCD.

## CRediT authorship contribution statement

**Shale A. Springer:** Conceptualization, Formal analysis, Investigation, Writing – Original Draft, Visualization. **Rishi Deshmukh:** Investigation. **Lambertus Klei:** Software, Formal analysis, Visualization. **Bernie Devlin:** Formal analysis, Writing – Review & Editing. **Jill R. Glausier:** Conceptualization, Project administration, Writing – Review & Editing. **David A. Lewis:** Conceptualization, Project administration, Writing – Review & Editing. **Susanne E. Ahmari:** Conceptualization, Supervision, Writing – Review & Editing, Funding acquisition.

## Acknowledgments

The authors would also like to thank the patients and their families for their generous tissue donation. Tissue from some subjects was obtained from the NIH NeuroBioBank at the University of Pittsburgh Brain Tissue Donation Program. We would also like to thank Kelly Rogers for her assistance with tissue blocking and cutting, as well as Dr Sue Johnston for her assistance with clinical assessments of post-mortem subjects. We would like to thank Dr Marianne Seney for conceptual feedback and use of laboratory equipment. This project utilized the services of the University of Pittsburgh Health Sciences Sequencing Core at UPMC Children’s Hospital of Pittsburgh for RNAseq library generation from RNA.

## Data Availability

RNA sequencing data will be made available in the Gene Expression Omnibus (Accession number GSE272200). Quantitative PCR data is available upon request.

## References

Abdul, A. ur R.M., De Silva, B., Gary, R.K., 2017. The GSK3 kinase inhibitor lithium produces unexpected hyperphosphorylation of β-catenin, a GSK3 substrate, in human glioblastoma cells. Biol Open 7. 10.1242/bio.030874

Ballouz, S., Verleyen, W., Gillis, J., 2015. Guidance for RNA-seq co-expression network construction and analysis: safety in numbers. Bioinformatics 31, 2123–2130. 10.1093/bioinformatics/btv118

Bienvenu, O.J., Wang, Y., Shugart, Y.Y., Welch, J.M., Grados, M.A., Fyer, A.J., Rauch, S.L., McCracken, J.T., Rasmussen, S.A., Murphy, D.L., Cullen, B., Valle, D., Hoehn-Saric, R., Greenberg, B.D., Pinto, A., Knowles, J.A., Piacentini, J., Pauls, D.L., Liang, K.Y., Willour, V.L., Riddle, M., Samuels, J.F., Feng, G., Nestadt, G., 2009. Sapap3 and pathological grooming in humans: Results from the OCD collaborative genetics study. American Journal of Medical Genetics Part B: Neuropsychiatric Genetics 150B, 710–720. 10.1002/ajmg.b.30897

Biria, M., Cantonas, L.-M., Banca, P., 2021. Magnetic Resonance Spectroscopy (MRS) and Positron Emission Tomography (PET) Imaging in Obsessive-Compulsive Disorder, in: Fineberg, N.A., Robbins, T.W. (Eds.), The Neurobiology and Treatment of OCD: Accelerating Progress, Current Topics in Behavioral Neurosciences. Springer International Publishing, Cham, pp. 231–268. 10.1007/7854_2020_201

Bour, L.J., Ackermans, L., Foncke, E.M.J., Cath, D., van der Linden, C., Visser Vandewalle, V., Tijssen, M.A., 2015. Tic related local field potentials in the thalamus and the effect of deep brain stimulation in Tourette syndrome: Report of three cases. Clinical Neurophysiology 126, 1578–1588. 10.1016/j.clinph.2014.10.217

Chakrabarty, K., Bhattacharyya, S., Christopher, R., Khanna, S., 2005. Glutamatergic Dysfunction in OCD. Neuropsychopharmacol 30, 1735–1740. 10.1038/sj.npp.1300733

Cope, D.W., Hughes, S.W., Crunelli, V., 2005. GABAA Receptor-Mediated Tonic Inhibition in Thalamic Neurons. J. Neurosci. 25, 11553–11563. 10.1523/JNEUROSCI.3362-05.2005

de Oliveira, K.C., Camilo, C., Gastaldi, V.D., Sant’Anna Feltrin, A., Lisboa, B.C.G., de Jesus Rodrigues de Paula, V., Moretto, A.C., Lafer, B., Hoexter, M.Q., Miguel, E.C., Maschietto, M., Brentani, H., 2021. Brain areas involved with obsessive-compulsive disorder present different DNA methylation modulation. BMC Genom Data 22, 45. 10.1186/s12863-021-00993-0

de Wit, S.J., de Vries, F.E., van der Werf, Y.D., Cath, D.C., Heslenfeld, D.J., Veltman, E.M., van Balkom, A.J.L.M., Veltman, D.J., van den Heuvel, O.A., 2012. Presupplementary Motor Area Hyperactivity During Response Inhibition: A Candidate Endophenotype of Obsessive-Compulsive Disorder. AJP 169, 1100–1108. 10.1176/appi.ajp.2012.12010073

Dolga, A.M., Letsche, T., Gold, M., Doti, N., Bacher, M., Chiamvimonvat, N., Dodel, R., Culmsee, C., 2012. Activation of KCNN3/SK3/KCa2.3 channels attenuates enhanced calcium influx and inflammatory cytokine production in activated microglia. Glia 60, 2050–2064. 10.1002/glia.22419

Fromer, M., Roussos, P., Sieberts, S.K., Johnson, J.S., Kavanagh, D.H., Perumal, T.M., Ruderfer, D.M., Oh, E.C., Topol, A., Shah, H.R., Klei, L.L., Kramer, R., Pinto, D., Gümüş, Z.H., Cicek, A.E., Dang, K.K., Browne, A., Lu, C., Xie, L., Readhead, B., Stahl, E.A., Xiao, J., Parvizi, M., Hamamsy, T., Fullard, J.F., Wang, Y.-C., Mahajan, M.C., Derry, J.M.J., Dudley, J.T., Hemby, S.E., Logsdon, B.A., Talbot, K., Raj, T., Bennett, D.A., De Jager, P.L., Zhu, J., Zhang, B., Sullivan, P.F., Chess, A., Purcell, S.M., Shinobu, L.A., Mangravite, L.M., Toyoshiba, H., Gur, R.E., Hahn, C.-G., Lewis, D.A., Haroutunian, V., Peters, M.A., Lipska, B.K., Buxbaum, J.D., Schadt, E.E., Hirai, K., Roeder, K., Brennand, K.J., Katsanis, N., Domenici, E., Devlin, B., Sklar, P., 2016. Gene expression elucidates functional impact of polygenic risk for schizophrenia. Nat Neurosci 19, 1442–1453. 10.1038/nn.4399

Gerentes, M., Pelissolo, A., Rajagopal, K., Tamouza, R., Hamdani, N., 2019. Obsessive-Compulsive Disorder: Autoimmunity and Neuroinflammation. Curr Psychiatry Rep 21, 78. 10.1007/s11920-019-1062-8

Goodman, W.K., Storch, E.A., Sheth, S.A., 2021. Harmonizing the Neurobiology and Treatment of Obsessive-Compulsive Disorder. Am J Psychiatry 178, 17–29. 10.1176/appi.ajp.2020.20111601

Grünblatt, E., 2021. Genetics of OCD and Related Disorders; Searching for Shared Factors, in: Fineberg, N.A., Robbins, T.W. (Eds.), The Neurobiology and Treatment of OCD: Accelerating Progress, Current Topics in Behavioral Neurosciences. Springer International Publishing, Cham, pp. 1–16. 10.1007/7854_2020_194

Gürsel, D.A., Avram, M., Sorg, C., Brandl, F., Koch, K., 2018. Frontoparietal areas link impairments of large-scale intrinsic brain networks with aberrant fronto-striatal interactions in OCD: a meta-analysis of resting-state functional connectivity. Neurosci Biobehav Rev 87, 151–160. 10.1016/j.neubiorev.2018.01.016

Hazari, N., Narayanaswamy, J.C., Venkatasubramanian, G., 2019. Neuroimaging findings in obsessive–compulsive disorder: A narrative review to elucidate neurobiological underpinnings. Indian J Psychiatry 61, S9–S29. 10.4103/psychiatry.IndianJPsychiatry_525_18

Huys, D., Bartsch, C., Koester, P., Lenartz, D., Maarouf, M., Daumann, J., Mai, J.K., Klosterkötter, J., Hunsche, S., Visser-Vandewalle, V., Woopen, C., Timmermann, L., Sturm, V., Kuhn, J., 2016. Motor Improvement and Emotional Stabilization in Patients With Tourette Syndrome After Deep Brain Stimulation of the Ventral Anterior and Ventrolateral Motor Part of the Thalamus. Biological Psychiatry, Extrapyramidal Motor Syndromes and the Striatum 79, 392–401. 10.1016/j.biopsych.2014.05.014

Jin, W., Sugaya, A., Tsuda, T., Ohguchi, H., Sugaya, E., 2000. Relationship between large conductance calcium-activated potassium channel and bursting activity. Brain Res 860, 21–28. 10.1016/s0006-8993(00)01943-0

Jung, W.H., Yücel, M., Yun, J.-Y., Yoon, Y.B., Cho, K.I.K., Parkes, L., Kim, S.N., Kwon, J.S., 2017. Altered functional network architecture in orbitofronto-striato-thalamic circuit of unmedicated patients with obsessive-compulsive disorder. Human Brain Mapping 38, 109–119. 10.1002/hbm.23347

Karolewicz, B., Maciag, D., O’Dwyer, G., Stockmeier, C.A., Feyissa, A.M., Rajkowska, G., 2010. Reduced level of glutamic acid decarboxylase-67 kDa in the prefrontal cortex in major depression. International Journal of Neuropsychopharmacology 13, 411–420. 10.1017/S1461145709990587

Kasten, C.R., Holmgren, E.B., Wills, T.A., 2019. Metabotropic Glutamate Receptor Subtype 5 in Alcohol-Induced Negative Affect. Brain Sci 9. 10.3390/brainsci9080183

Kasten, M.R., Rudy, B., Anderson, M.P., 2007. Differential regulation of action potential firing in adult murine thalamocortical neurons by Kv3.2, Kv1, and SK potassium and N-type calcium channels. J Physiol 584, 565–582. 10.1113/jphysiol.2007.141135

Khurshid, K.A., 2020. High frequency repetitive transcranial magnetic stimulation of supplementary motor cortex for obsessive compulsive disorder. Medical Hypotheses 137, 109529. 10.1016/j.mehy.2019.109529

Kim, D., Song, I., Keum, S., Lee, T., Jeong, M.-J., Kim, S.-S., McEnery, M.W., Shin, H.-S., 2001. Lack of the Burst Firing of Thalamocortical Relay Neurons and Resistance to Absence Seizures in Mice Lacking α1G T-Type Ca2+ Channels. Neuron 31, 35–45. 10.1016/S0896-6273(01)00343-9

Kim, T., Kim, M., Jung, W.H., Kwak, Y.B., Moon, S.-Y., Kyungjin Lho, S., Lee, J., Kwon, J.S., 2022. Unbalanced fronto-pallidal neurocircuit underlying set shifting in obsessive-compulsive disorder. Brain 145, 979–990. 10.1093/brain/awab483

Langfelder, P., Horvath, S., 2008. WGCNA: an R package for weighted correlation network analysis. BMC Bioinformatics 9, 559. 10.1186/1471-2105-9-559

Langfelder, P., Horvath, S., 2007. Eigengene networks for studying the relationships between co-expression modules. BMC Syst Biol 1, 54. 10.1186/1752-0509-1-54

Langfelder, P., Zhang, B., Horvath, S., 2008. Defining clusters from a hierarchical cluster tree: the Dynamic Tree Cut package for R. Bioinformatics 24, 719–720. 10.1093/bioinformatics/btm563

Law, C.W., Chen, Y., Shi, W., Smyth, G.K., 2014. voom: precision weights unlock linear model analysis tools for RNA-seq read counts. Genome Biol 15, R29. 10.1186/gb-2014-15-2-r29

Leek, J.T., Johnson, W.E., Parker, H.S., Jaffe, A.E., Storey, J.D., 2012. The sva package for removing batch effects and other unwanted variation in high-throughput experiments. Bioinformatics 28, 882–883. 10.1093/bioinformatics/bts034

Li, K., Fan, L., Cui, Y., Wei, X., He, Y., Yang, J., Lu, Y., Li, W., Shi, W., Cao, L., Cheng, L., Li, A., You, B., Jiang, T., 2022. The human mediodorsal thalamus: Organization, connectivity, and function. NeuroImage 249, 118876. 10.1016/j.neuroimage.2022.118876

Li, K., Zhang, Haisan, Yang, Y., Zhu, J., Wang, B., Shi, Y., Li, X., Meng, Z., Lv, L., Zhang, Hongxing, 2019. Abnormal functional network of the thalamic subregions in adult patients with obsessive-compulsive disorder. Behavioural Brain Research 371, 111982. 10.1016/j.bbr.2019.111982

Liao, Y., Wang, J., Jaehnig, E.J., Shi, Z., Zhang, B., 2019. WebGestalt 2019: gene set analysis toolkit with revamped UIs and APIs. Nucleic Acids Research 47, W199–W205. 10.1093/nar/gkz401

Lisboa, B.C.G., Oliveira, K.C., Tahira, A.C., Barbosa, A.R., Feltrin, A.S., Gouveia, G., Lima, L., Feio dos Santos, A.C., Martins, D.C., Puga, R.D., Moretto, A.C., De Bragança Pereira, C.A., Lafer, B., Leite, R.E.P., Ferretti-Rebustini, R.E.D.L., Farfel, J.M., Grinberg, L.T., Jacob-Filho, W., Miguel, E.C., Hoexter, M.Q., Brentani, H., 2019. Initial findings of striatum tripartite model in OCD brain samples based on transcriptome analysis. Sci Rep 9, 3086. 10.1038/s41598-019-38965-1

Lv, Q., Wang, Zhen, Zhang, C., Fan, Q., Zhao, Q., Zeljic, K., Sun, B., Xiao, Z., Wang, Zheng, 2017. Divergent Structural Responses to Pharmacological Interventions in Orbitofronto-Striato-Thalamic and Premotor Circuits in Obsessive-Compulsive Disorder. EBioMedicine 22, 242–248. 10.1016/j.ebiom.2017.07.021

Macy, A.S., Theo, J.N., Kaufmann, S.C.V., Ghazzaoui, R.B., Pawlowski, P.A., Fakhry, H.I., Cassmassi, B.J., IsHak, W.W., 2013. Quality of life in obsessive compulsive disorder. CNS Spectrums 18, 21–33. 10.1017/S1092852912000697

Mahjani, B., Bey, K., Boberg, J., Burton, C., 2021. Genetics of obsessive-compulsive disorder. Psychol Med 51, 2247–2259. 10.1017/S0033291721001744

Mai, J.K., Majtanik, M., Paxinos, G., 2015. Atlas of the Human Brain. Academic Press.

Marcks, B.A., Weisberg, R.B., Dyck, I., Keller, M.B., 2011. Longitudinal course of obsessive-compulsive disorder in patients with anxiety disorders: a 15-year prospective follow-up study. Compr Psychiatry 52, 670–677. 10.1016/j.comppsych.2011.01.001

McFarland, N.R., Haber, S.N., 2002. Thalamic Relay Nuclei of the Basal Ganglia Form Both Reciprocal and Nonreciprocal Cortical Connections, Linking Multiple Frontal Cortical Areas. J Neurosci 22, 8117–8132. 10.1523/JNEUROSCI.22-18-08117.2002

McFarland, N.R., Haber, S.N., 2000. Convergent Inputs from Thalamic Motor Nuclei and Frontal Cortical Areas to the Dorsal Striatum in the Primate. J. Neurosci. 20, 3798–3813. 10.1523/JNEUROSCI.20-10-03798.2000

Mitchell, A.S., Chakraborty, S., 2013. What does the mediodorsal thalamus do? Front Syst Neurosci 7, 37. 10.3389/fnsys.2013.00037

Norman, L.J., Taylor, S.F., Liu, Y., Radua, J., Chye, Y., De Wit, S.J., Huyser, C., Karahanoglu, F.I., Luks, T., Manoach, D., Mathews, C., Rubia, K., Suo, C., van den Heuvel, O.A., Yücel, M., Fitzgerald, K., 2019. Error Processing and Inhibitory Control in Obsessive-Compulsive Disorder: A Meta-analysis Using Statistical Parametric Maps. Biological Psychiatry, Neurodevelopmental Mechanisms of Obsessive-Compulsive Disorder and Autism 85, 713–725. 10.1016/j.biopsych.2018.11.010

Paydar, A., Lee, B., Gangadharan, G., Lee, S., Hwang, E.M., Shin, H.-S., 2014. Extrasynaptic GABAA receptors in mediodorsal thalamic nucleus modulate fear extinction learning. Molecular Brain 7, 39. 10.1186/1756-6606-7-39

Perera, M.P.N., Gotsis, E.S., Bailey, N.W., Fitzgibbon, B.M., Fitzgerald, P.B., 2024. Exploring functional connectivity in large-scale brain networks in obsessive-compulsive disorder: a systematic review of EEG and fMRI studies. Cereb Cortex 34, bhae327. 10.1093/cercor/bhae327

Perera, M.P.N., Mallawaarachchi, S., Bailey, N.W., Murphy, O.W., Fitzgerald, P.B., 2023. Obsessive-compulsive disorder (OCD) is associated with increased electroencephalographic (EEG) delta and theta oscillatory power but reduced delta connectivity. Journal of Psychiatric Research 163, 310–317. 10.1016/j.jpsychires.2023.05.026

Piantadosi, S.C., Chamberlain, B.L., Glausier, J.R., Lewis, D.A., Ahmari, S.E., 2019. Lower excitatory synaptic gene expression in orbitofrontal cortex and striatum in an initial study of subjects with obsessive compulsive disorder. Molecular Psychiatry 1. 10.1038/s41380-019-0431-3

Piantadosi, S.C., McClain, L.L., Klei, L., Wang, J., Chamberlain, B.L., Springer, S.A., Lewis, D.A., Devlin, B., Ahmari, S.E., 2021. Transcriptome alterations are enriched for synapse-associated genes in the striatum of subjects with obsessive-compulsive disorder. Transl Psychiatry 11, 1–11. 10.1038/s41398-021-01290-1

Priori, A., Giannicola, G., Rosa, M., Marceglia, S., Servello, D., Sassi, M., Porta, M., 2013. Deep brain electrophysiological recordings provide clues to the pathophysiology of Tourette syndrome. Neuroscience & Biobehavioral Reviews, The multifaceted nature of Tourette syndrome: Pre-clinical, clinical and therapeutic issues 37, 1063–1068. 10.1016/j.neubiorev.2013.01.011

Ritchie, M.E., Phipson, B., Wu, D., Hu, Y., Law, C.W., Shi, W., Smyth, G.K., 2015. limma powers differential expression analyses for RNA-sequencing and microarray studies. Nucleic Acids Res 43, e47. 10.1093/nar/gkv007

Rotge, J.Y., Aouizerate, B., Amestoy, V., Lambrecq, V., Langbour, N., Nguyen, T.H., Dovero, S., Cardoit, L., Tignol, J., Bioulac, B., Burbaud, P., Guehl, D., 2012. The associative and limbic thalamus in the pathophysiology of obsessive-compulsive disorder: an experimental study in the monkey. Transl Psychiatry 2, e161. 10.1038/tp.2012.88

Rotge, J.-Y., Guehl, D., Dilharreguy, B., Cuny, E., Tignol, J., Bioulac, B., Allard, M., Burbaud, P., Aouizerate, B., 2008. Provocation of obsessive–compulsive symptoms: a quantitative voxel-based meta-analysis of functional neuroimaging studies. J Psychiatry Neurosci 33, 405–412.

Ruscio, A.M., Stein, D.J., Chiu, W.T., Kessler, R.C., 2010. The Epidemiology of Obsessive-Compulsive Disorder in the National Comorbidity Survey Replication. Mol Psychiatry 15, 53–63. 10.1038/mp.2008.94

Shao, J., Liu, Y., Gao, D., Tu, J., Yang, F., 2021. Neural Burst Firing and Its Roles in Mental and Neurological Disorders. Front. Cell. Neurosci. 15. 10.3389/fncel.2021.741292

Sharma, E., Thennarasu, K., Reddy, Y.C.J., 2014. Long-Term Outcome of Obsessive-Compulsive Disorder in Adults: A Meta-Analysis. J Clin Psychiatry 75, 10746. 10.4088/JCP.13r08849

Shephard, E., Batistuzzo, M.C., Hoexter, M.Q., Stern, E.R., Zuccolo, P.F., Ogawa, C.Y., Silva, R.M., Brunoni, A.R., Costa, D.L., Doretto, V., Saraiva, L., Cappi, C., Shavitt, R.G., Simpson, H.B., van den Heuvel, O.A., Miguel, E.C., 2021. Neurocircuit models of obsessive-compulsive disorder: limitations and future directions for research. Braz J Psychiatry 44, 187–200. 10.1590/1516-4446-2020-1709

Sherman, S.M., 2004. Interneurons and triadic circuitry of the thalamus. Trends in Neurosciences 27, 670–675. 10.1016/j.tins.2004.08.003

Shmelkov, S.V., Hormigo, A., Jing, D., Proenca, C.C., Bath, K.G., Milde, T., Shmelkov, E., Kushner, J.S., Baljevic, M., Dincheva, I., Murphy, A.J., Valenzuela, D.M., Gale, N.W., Yancopoulos, G.D., Ninan, I., Lee, F.S., Rafii, S., 2010. Slitrk5 deficiency impairs corticostriatal circuitry and leads to obsessive-compulsive–like behaviors in mice. Nat Med 16, 598–602. 10.1038/nm.2125

Siddiqui, T., Lively, S., Ferreira, R., Wong, R., Schlichter, L.C., 2014. Expression and Contributions of TRPM7 and KCa2.3/SK3 Channels to the Increased Migration and Invasion of Microglia in Anti-Inflammatory Activation States. PLOS ONE 9, e106087. 10.1371/journal.pone.0106087

Strom, N.I., Burton, C.L., Iyegbe, C., Silzer, T., Antonyan, L., Pool, R., Lemire, M., Crowley, J.J., Hottenga, J.-J., Ivanov, V.Z., Larsson, H., Lichtenstein, P., Magnusson, P., Rück, C., Schachar, R., Wu, H.M., Cath, D., Crosbie, J., Mataix-Cols, D., Boomsma, D.I., Mattheisen, M., Meier, S.M., Smit, D.J.A., Arnold, P.D., 2024. Genome-Wide Association Study of Obsessive-Compulsive Symptoms including 33,943 individuals from the general population. Mol Psychiatry 1–10. 10.1038/s41380-024-02489-6

Thompson, M., Weickert, C.S., Wyatt, E., Webster, M.J., 2009. Decreased glutamic acid decarboxylase67 mRNA expression in multiple brain areas of patients with schizophrenia and mood disorders. Journal of Psychiatric Research 43, 970–977. 10.1016/j.jpsychires.2009.02.005

Welch, J.M., Lu, J., Rodriguiz, R.M., Trotta, N.C., Peca, J., Ding, J.-D., Feliciano, C., Chen, M., Adams, J.P., Luo, J., Dudek, S.M., Weinberg, R.J., Calakos, N., Wetsel, W.C., Feng, G., 2007. Cortico-striatal synaptic defects and OCD-like behaviors in SAPAP3 mutant mice. Nature 448, 894–900. 10.1038/nature06104

White, K., Yang, P., Li, L., Farshori, A., Medina, A.E., Zielke, H.R., 2018. Effect of Postmortem Interval and Years in Storage on RNA Quality of Tissue at a Repository of the NIH NeuroBioBank. Biopreservation and Biobanking 16, 148–157. 10.1089/bio.2017.0099

Zhang, B., Horvath, S., 2005. A General Framework for Weighted Gene Co-Expression Network Analysis. Statistical Applications in Genetics and Molecular Biology 4. 10.2202/1544-6115.1128

Zobeiri, M., Chaudhary, R., Blaich, A., Rottmann, M., Herrmann, S., Meuth, P., Bista, P., Kanyshkova, T., Lüttjohann, A., Narayanan, V., Hundehege, P., Meuth, S.G., Romanelli, M.N., Urbano, F.J., Pape, H.-C., Budde, T., Ludwig, A., 2019. The Hyperpolarization-Activated HCN4 Channel is Important for Proper Maintenance of Oscillatory Activity in the Thalamocortical System. Cereb Cortex 29, 2291–2304. 10.1093/cercor/bhz047

