## Supplemental Tables 1-10 for "Transcriptomes of higher order thalamic nuclei in obsessive compulsive disorder": Supplemental Table 1.pdf

**Unaffected Comparison Subjects**

| Pair | Case | Sex | Race | Handedness | Age | BMI | PMI | pH[a] | Subject RIN | Tissue Storage Time (mo)[b] | Medications ATOD[c] | Tobacco ATOD | Manner of Death | Cause of Death |
|---|---|---|---|---|---|---|---|---|---|---|---|---|---|---|
| 1 | 1159 | M | W | R | 51 | 38.5 | 16.7 | 6.5 | 7.6 | 201.8 | O | N | Natural | Cardiovascular Disease |
| 2 | 1293 | F | W | R | 65 | 19.7 | 18.5 | 6.5 | 7.0 | 182.6 | N | N | Accidental | Blunt Force Trauma |
| 3 | 13032 | M | W | R | 52 | 29.3 | 9.0 | 6.3 | 8.2 | 81.2 | N | Y | Natural | Pulmonary Embolism |
| 4 | 1092 | F | B | R | 40 | 39.3 | 16.6 | 6.7 | 8.0 | 209.1 | O | N | Natural | Mitral Valve Prolapse |
| 5 | 1543 | F | W | R | 45 | 23.0 | 17.9 | 6.8 | 7.4 | 136.2 | O | Y | Natural | Subarachnoid Hemorrhage |
| 6 | 1391 | F | W | L | 51 | 28.3 | 7.8 | 6.5 | 7.1 | 165.5 | O | Y | Natural | Cardiovascular Disease |
| 7 | 1083 | M | W | R | 20 | 27.2 | 19.9 | 6.6 | 8.8 | 210.2 | N | Y | Accidental | Blunt Force Trauma |
| 8 | 13237 | M | W | R | 25 | 22.7 | 22.3 | 6.2 | 8.2 | 53.3 | N | Y | Accidental | Blunt Force Trauma |
| 9 | 1637 | M | W | L | 46 | 32.9 | 16.6 | 6.9 | 8.2 | 121.7 | N | N | Natural | Cardiovascular Disease |
| 10 | 1282 | F | W | R | 39 | 30.6 | 24.5 | 6.8 | 7.5 | 184.9 | N | N | Natural | Cardiovascular Disease |
| 11 | 1834 | F | B | R | 39 | 41.0 | 13.5 | 6.8 | 8.2 | 86.0 | N | N | Undetermined | Undetermined |
| | **Mean** | | | | **43** | **30.2** | **16.7** | **6.6** | **7.8** | **148.4** | | | | |
| | **Standard Deviation** | | | | **12.6** | **7.1** | **5.1** | **0.2** | **0.6** | **56.1** | | | | |
| | **Counts** | **5M/6F** | **9W/2B** | **9R/2L** | | | | | | | | **5Y/6N** | | |

**Obsessive-Compulsive Disorder Subjects**

| Pair | Case | DSM-IV Obsessive-Compulsive Disorder Diagnosis ATOD | Duration of Obsessive-Compulsive Disorder Diagnosis | DSM-IV Co-Morbid Substance Use Diagnosis ATOD | Sex | Race | Handedness | Age | BMI | PMI | pH[a] | Subject RIN | Tissue Storage Time (mo)[b] | Medications ATOD[c] | Tobacco ATOD | Manner of Death | Cause of Death |
|---|---|---|---|---|---|---|---|---|---|---|---|---|---|---|---|---|---|
| 1 | 1413 | Obsessive-Compulsive Disorder | 46 | None | M | W | R | 52 | 34.8 | 17.4 | 6.5 | 8.0 | 160.4 | D P | N | Suicide | Fluvoxamine Overdose |
| 2 | 1424^ | Obsessive-Compulsive Disorder | U | None | F | W | R | 69 | 39.3 | 10.5 | 6.7 | 7.1 | 158.5 | O | N | Natural | Cardiomyopathy |
| 3 | 1435 | Obsessive-Compulsive Disorder | 10 | None | M | W | L | 52 | 26.0 | 7.9 | 6.7 | 8.0 | 157.3 | N | N | Natural | Cardiovascular Disease |
| 4 | 1505 | Obsessive-Compulsive Disorder | 34 | None | F | W | L | 42 | 19.3 | 23.5 | 6.4 | 8.2 | 144.9 | B C D O P | N | Natural | Cardiovascular Disease |
| 5 | 1785* | Obsessive-Compulsive Disorder | 12 | Alcohol Dependence; Sedative or Hypnotic or Anxiolytic Dependence | F | W | R | 50 | 23.1 | 29.8 | 6.9 | 7.3 | 94.9 | B C D P | Y | Suicide | Drowning |
| 6 | 13051 | Obsessive-Compulsive Disorder | 1 | None | F | W | R | 47 | 23.8 | 10.8 | 6.6 | 7.6 | 78.2 | B C D O | Y | Natural | Valvular Heart Disease |
| 7 | 13022 | Obsessive-Compulsive Disorder | 14 | None | M | W | R | 20 | 23.6 | 24.4 | 6.8 | 8.2 | 82.3 | O | Y | Suicide | Hydrogen Sulfide Overdose |
| 8 | 13173 | Obsessive-Compulsive Disorder | 22 | Sedative or Hypnotic or Anxiolytic Abuse | M | W | R | 30 | 30.0 | 21.0 | 6.5 | 8.6 | 61.0 | B D | N | Accidental | Combined Drug Overdose |
| 9 | 13125^ | Obsessive-Compulsive Disorder | 19 | Alcohol Dependence; Cannabis Abuse; Opioid Abuse | M | W | R | 49 | 34.9 | 14.9 | 6.4 | 8.2 | 67.4 | B C D O P | Y | Natural | Cardiovascular Disease |
| 10 | 13154 | Obsessive-Compulsive Disorder | 19 | Alcohol Dependence; Sedative or Hypnotic or Anxiolytic Dependence | F | W | R | 45 | 34.4 | 23.7 | 6.2 | 7.0 | 63.4 | B D O | N | Accidental | Blunt Force Trauma |
| 11 | 13169 | Obsessive-Compulsive Disorder | 11 | None | F | W | R | 41 | 21.5 | 8.7 | 6.7 | 8.7 | 61.5 | B D O | Y | Accidental | Blunt Force Trauma |
| | **Mean** | | **15.8** | | | | | **45.2** | **28.2** | **17.5** | **6.6** | **7.9** | **102.7** | | | | |
| | **Standard Deviation** | | **12.9** | | | | | **12.6** | **6.7** | **7.5** | **0.2** | **0.6** | **43.0** | | | | |
| | **Counts** | | | | **5M/6F** | **11W** | **9R/2L** | | | | | | | | **5Y/6N** | | |

Footnotes:

[a]Reported value is the mean of prefrontal and cerebellar pH values. Case 1083 is the exception with only the prefrontal value reported.
[b]Stored at -80C
[c]Medications at time of death: B, benzodiazepines; C, anticonvulsants; D, antidepressants; L, lithium; N, no medications; O, other medication(s); P, antipsychotic; U, unknown
*OCD in Remission ATOD
^1424: Cognitive Disorder NOS
^13125: Psychotic Disorder NOS

Abbreviations:

yr, years; BMI, body mass index; PMI, postmortem interval (hours); RIN, RNA integrity number; mo, months; ATOD, at time of death; F, female; M, male; B, Black; W, White; R, right; L, left; Y, yes; N, no; U, unknown; NOS, not otherwise specified
