## Supplemental Figures 1-7 for "Transcriptomes of higher order thalamic nuclei in obsessive compulsive disorder"

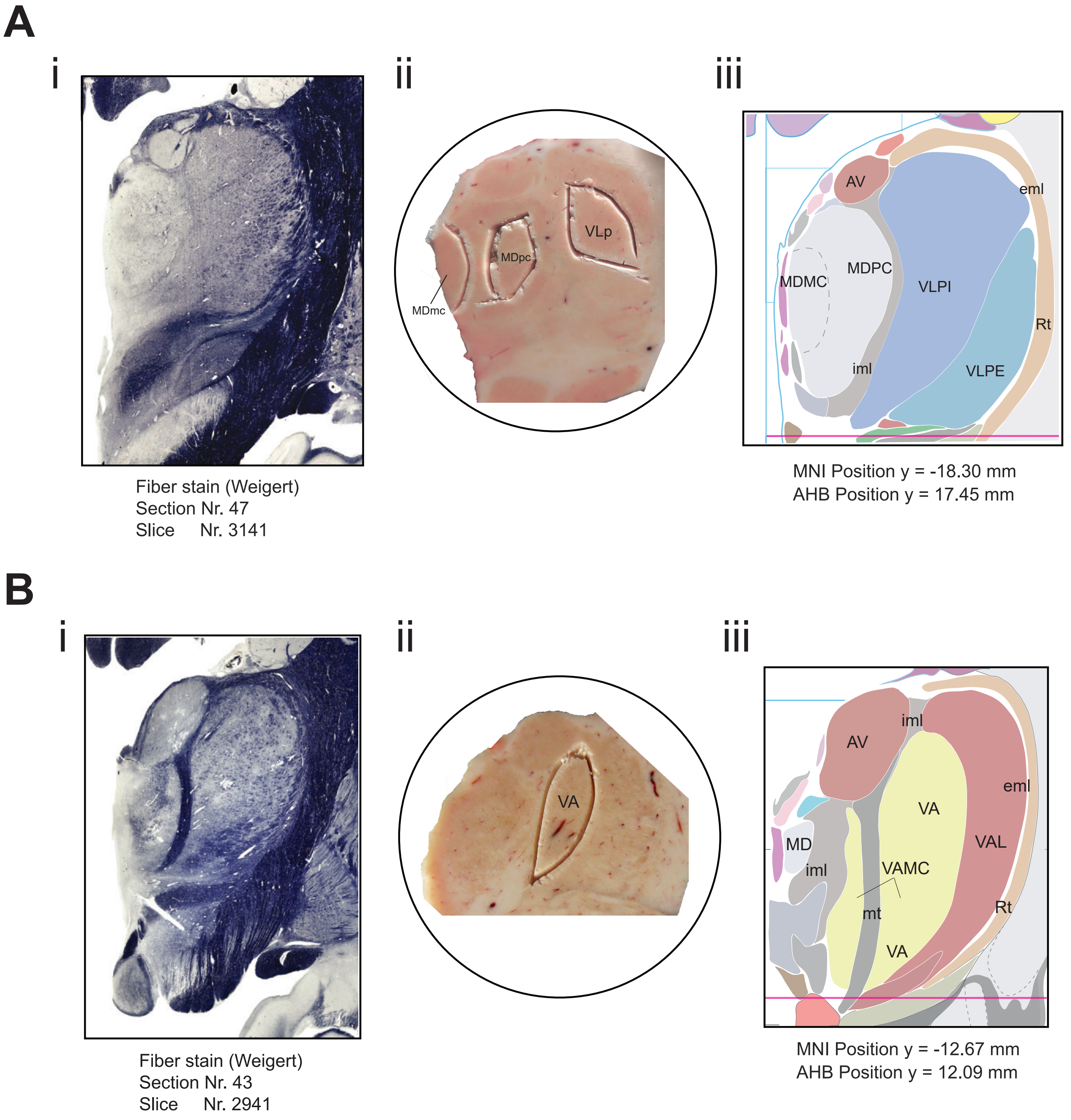


**Supplemental Figure 1.** Target collection areas. Fiber stain and atlas diagram figures adapted from 4th edition of the Atlas of the Human Brain (Mai, Majtanik and Paxinos 2016) under STM permission guidelines. A. (i) Fiber stain, (ii) representative tissue scoring for cryostat collection, and (iii) atlas diagram of collection areas for mediodorsal magnocellular (MDmc), mediodorsal parvocellular (MDpc) and posterior part of the ventrolateral (VLp) nuclei. Internal and external regions of VLp (VLPI and VLPE) were not differentiated during collection. B. (i) Fiber stain, (ii) representative tissue scoring, and (iii) atlas diagram of collection area for ventral anterior nucleus (VA). VA collection areas included VA magnocellular (VAMC) but not lateral VA (VAL). AV = anteroventral nucleus; eml = external medullary lamina; iml = internal medullary lamina; mt = mammillothalamic tract; Rt = thalamic reticular nucleus.

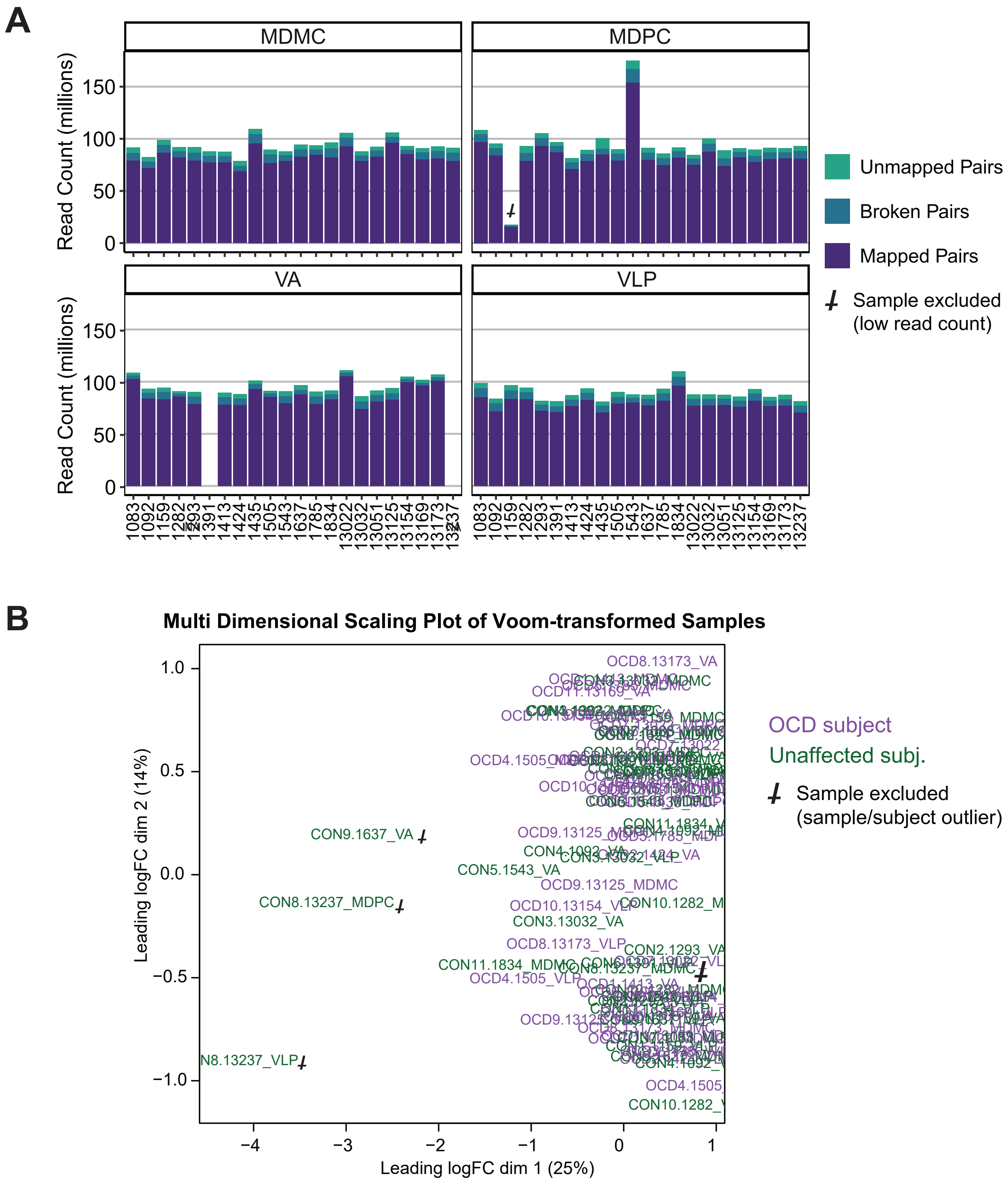


**Supplemental Figure 2. R**NA sequencing data quality assessment. A. RNA sequencing read counts by subject and region. Paired-end reads were mapped to the human reference sequence in intact pairs (purple) or in broken pairs (blue) or did not map (teal). Human subject number is identified along the X axis. B. Multidimensional scaling plot of voom-transformed transcript expression data for each sample. Three samples were identified as outliers and excluded from analysis, and the sole remaining sample from HU13237 was removed as well.

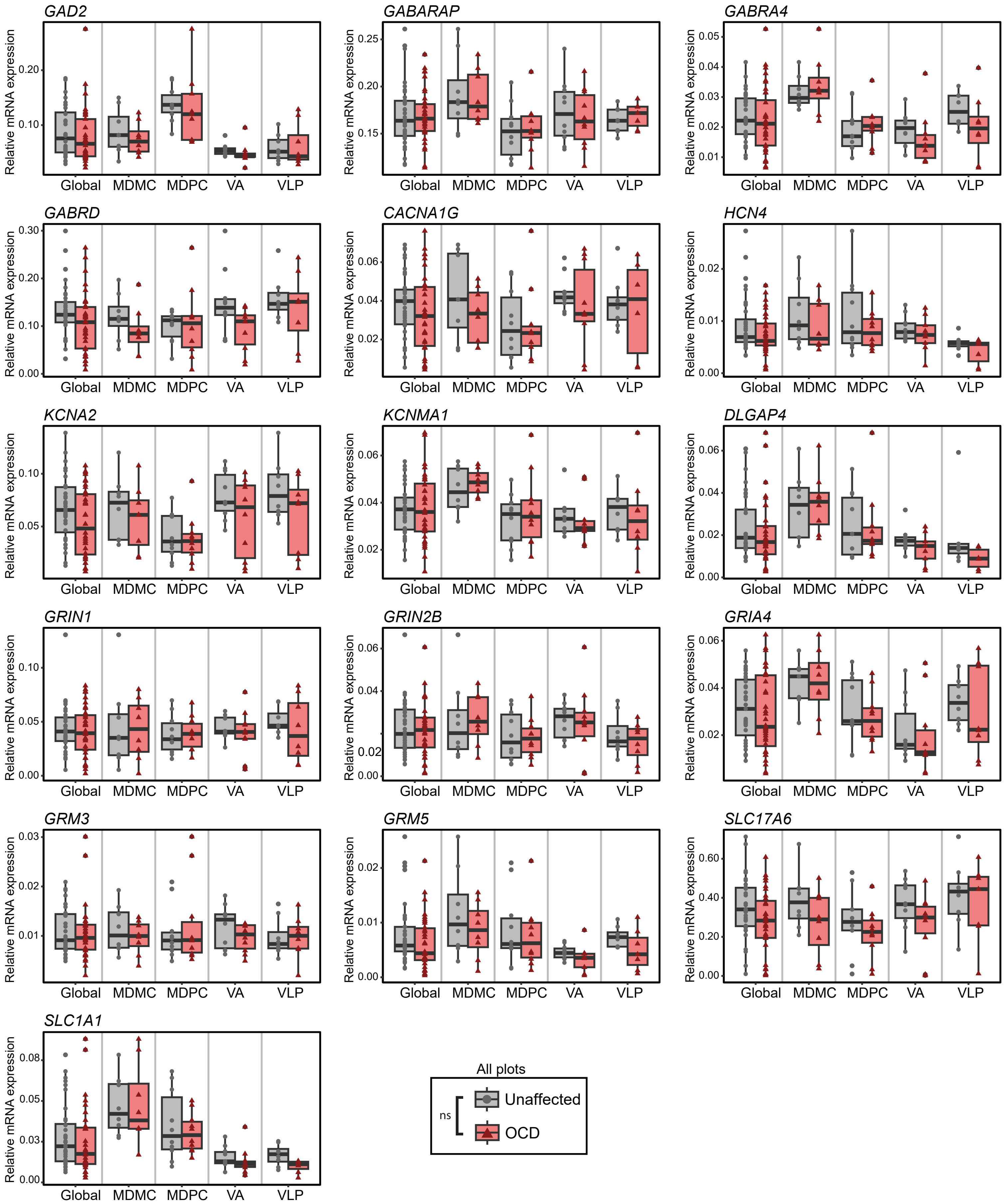


**Supplemental Figure 3.** Expression of additional candidate gene transcripts in thalamic nuclei of subjects with OCD and unaffected subjects. Boxplots depict relative expression levels of mRNA expression in each nucleus or across all nuclei (Global). ns = not significant. No significant effect of diagnosis or diagnosis-by-region interaction were identified.

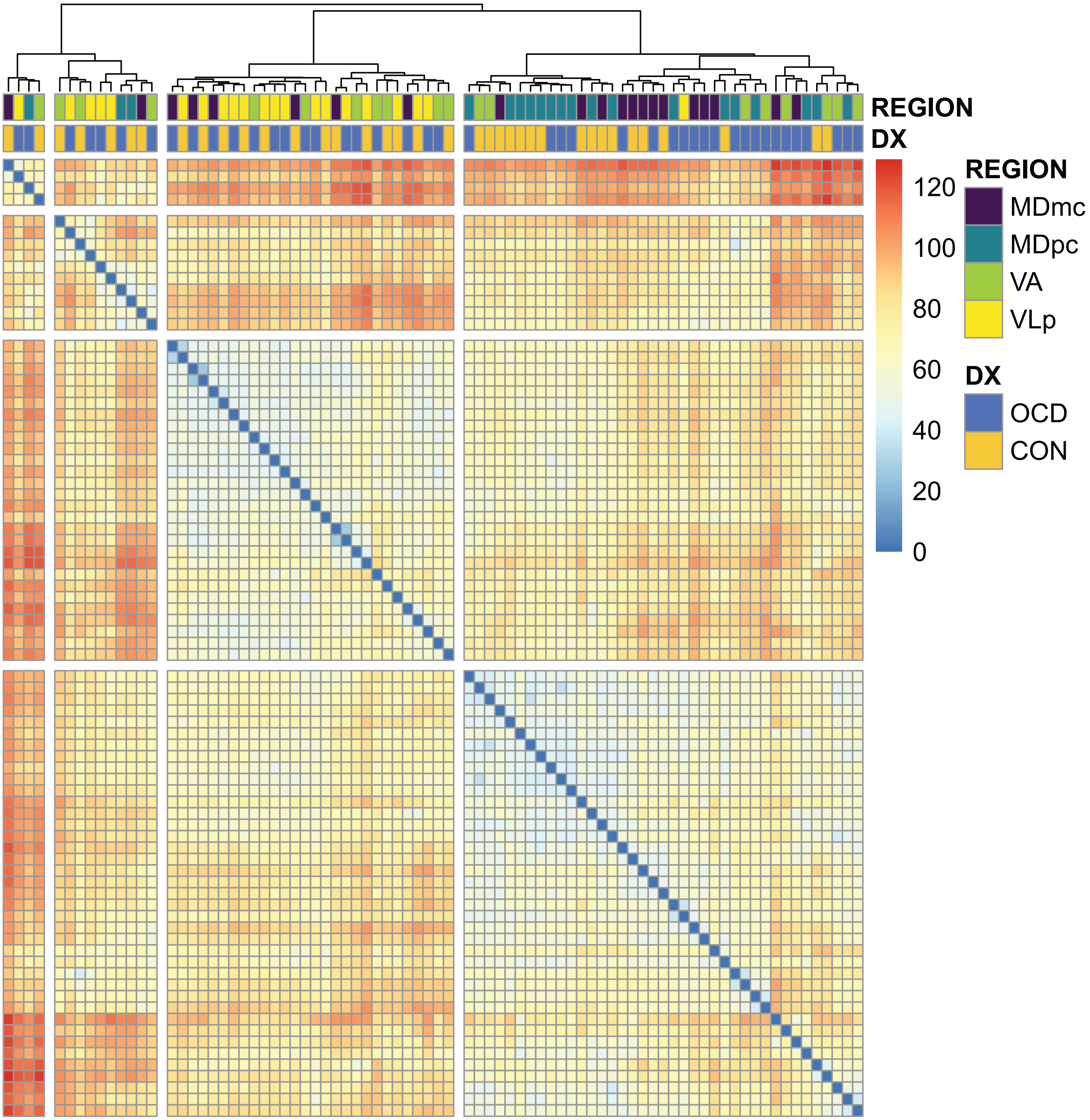


**Supplemental Figure 4.** Clustered dissimilarity heatmap of transcriptomic expression for all samples. Hierarchical clustering was performed using the Ward D2 method.

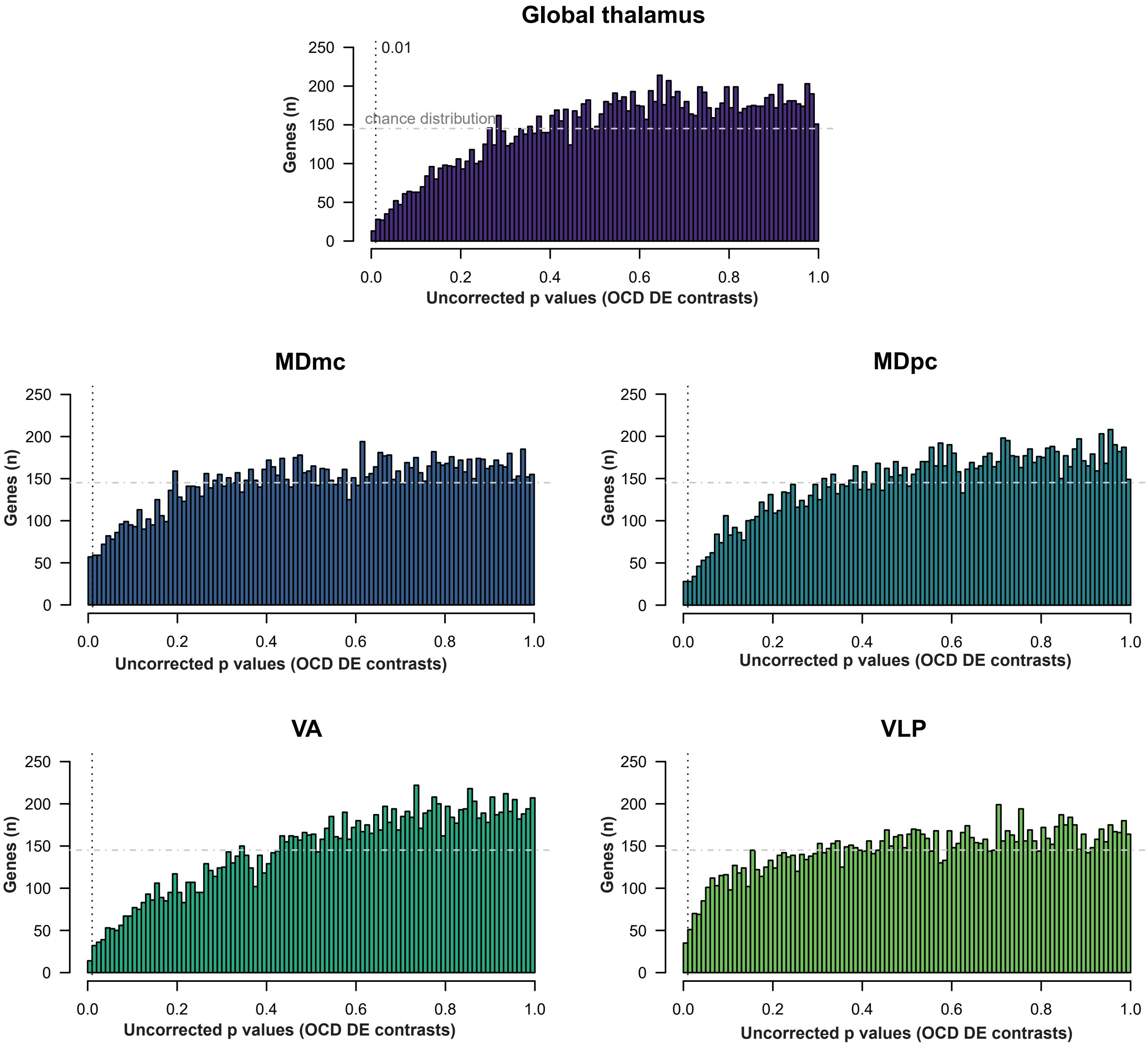


**Supplemental Figure 5.** Distribution of uncorrected p-values for OCD differential expression contrasts in global thalamic analysis or individual thalamic regions. The expected p-value distribution under the null hypothesis ("chance distribution") is indicated by a horizontal dash-dot line at y = 147, the expected height of all 1,000 histogram bins if every p value was equally likely. Observed leftward skew indicates fewer significant contrasts than expected by chance.

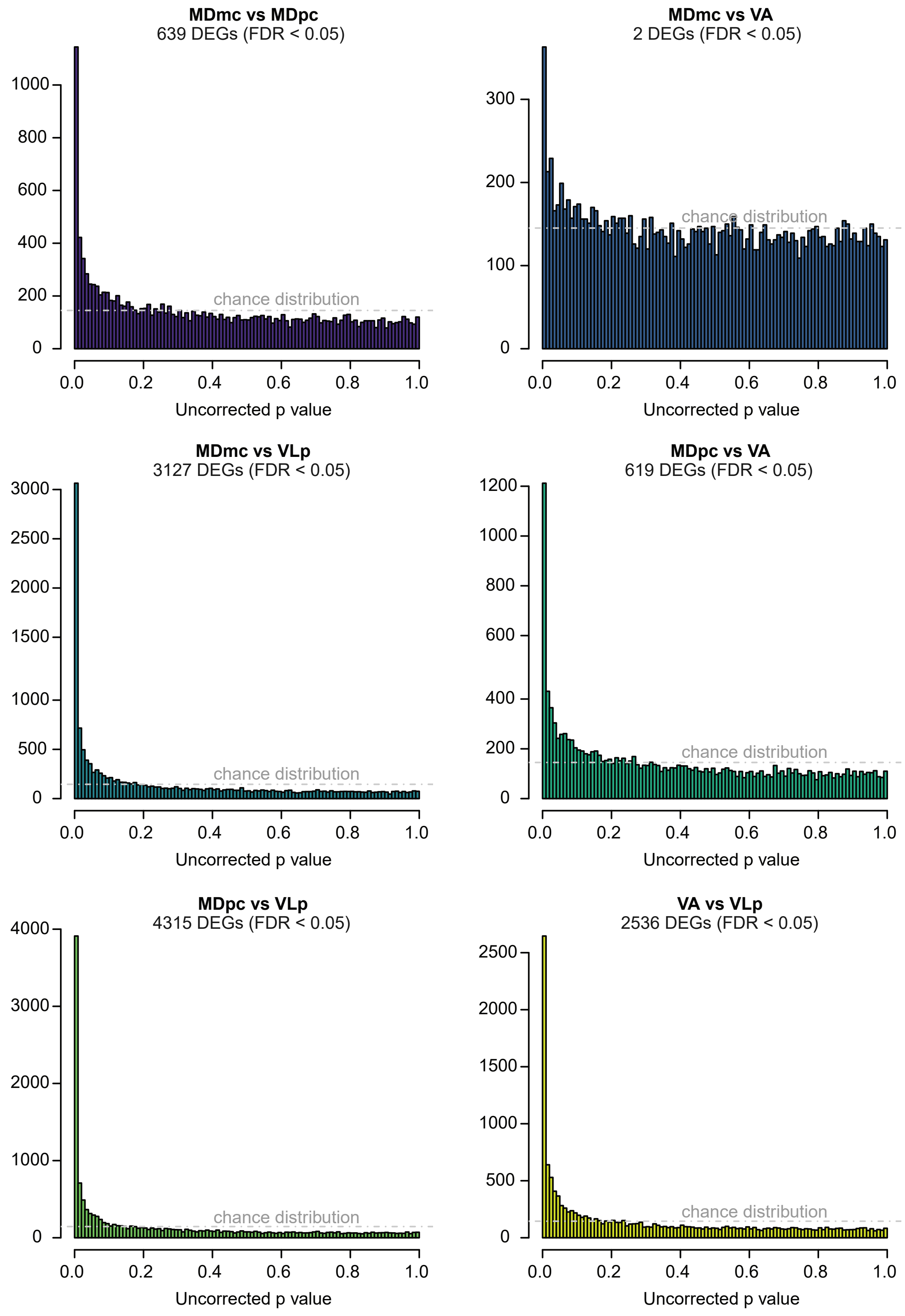


**Supplemental Figure 6.** P value distribution for differential gene expression between thalamic regions. The expected p-value distribution under the null hypothesis (“chance distribution”) is indicated by a horizontal dash-dot line at y =147, the expected height of all 1,000 histogram bins if every p value was equally likely. DEGs = differentially expressed genes; FDR = false discovery rate.

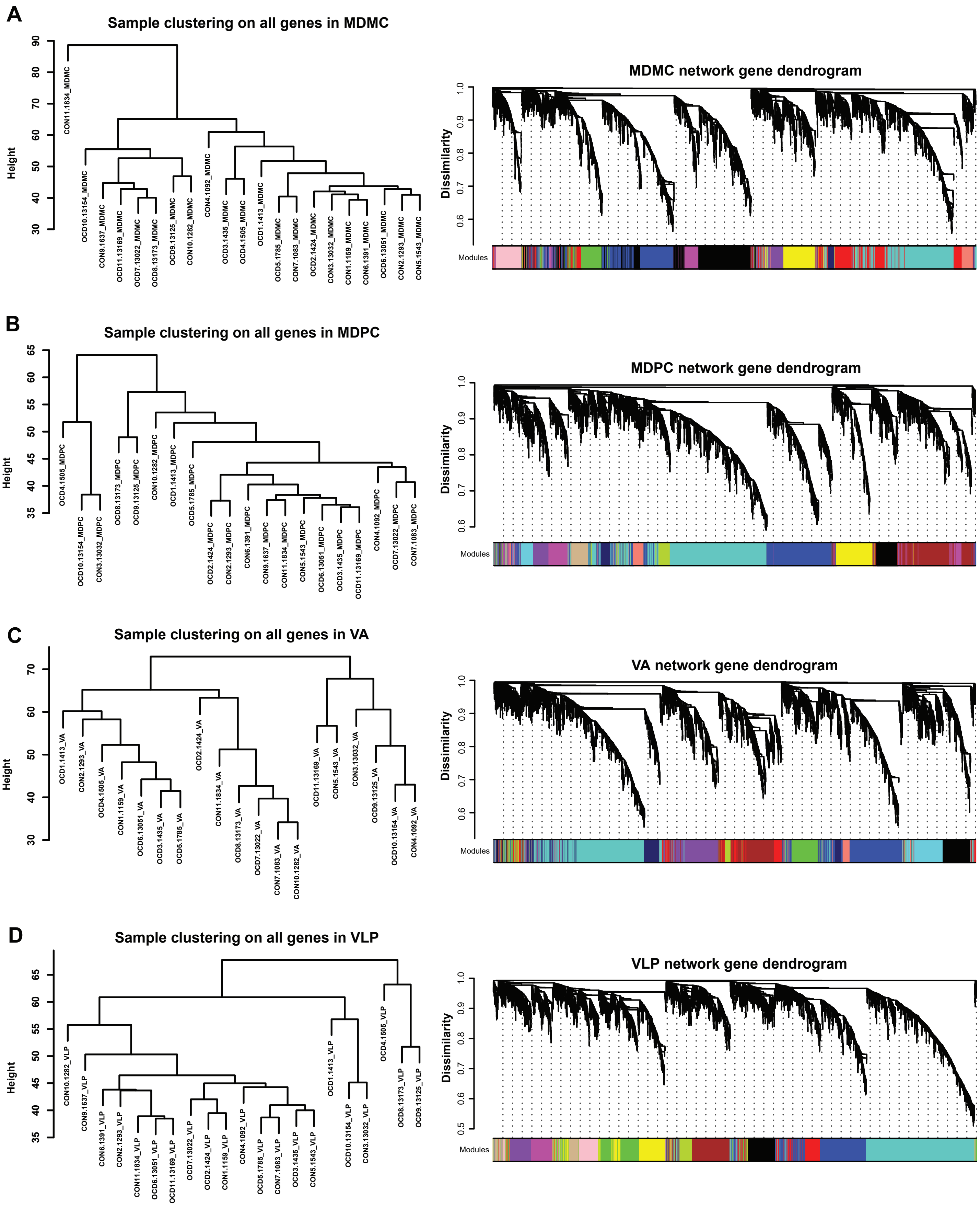


**Supplemental Figure 7.** Hierarchical sample clustering and weighted gene correlation network analysis (WGCNA) dendrograms for each thalamic nucleus (A – D). WGCNA networks were generated using biweight midcorrelation and signed-hybrid adjacency networks.
